# *Gnas* ablation in CD11c+ cells prevents high-fat diet-induced obesity by elevating adipose tissue catecholamine levels and thermogenesis

**DOI:** 10.1101/2022.01.27.478070

**Authors:** Liping Zeng, D. Scott Herdman, Jihyung Lee, Ailin Tao, Manasi Das, Samuel Bertin, Lars Eckmann, Sushil Mahata, Shwetha Devulapalli, Hemal H. Patel, Anthony J.A. Molina, Olivia Osborn, Maripat Corr, Eyal Raz, Nicholas J.G. Webster

## Abstract

CD11c+ immune cells are a potential therapeutic target for treatment of obesity-related insulin resistance and type 2 diabetes (T2D). In obesity, CD11c+ immune cells are recruited to white adipose tissue and create an inflammatory state that causes both insulin and catecholamine resistance. In this study, we found that ablation of *Gnas,* the gene that encodes Gas, in CD11c expressing cells protects mice from high-fat diet-induced obesity, glucose intolerance and insulin resistance. *Gnas^ΔCD11c^* mice (KO) had increased oxygen consumption, energy expenditure, and beigeing of white adipose tissue (WAT). Transplantation studies showed that the lean phenotype was conferred by bone marrow-derived cells and the absence of T and B cells by crossing the KO to a *Rag1^-^*^/-^ background did not alter the phenotype. Notably, we observed elevated norepinephrine and elevated cAMP signaling in the WAT of KO mice. The KO adipose tissue also had reduced expression of catecholamine transport and degradation enzymes. Collectively, our results identified an important role of Gas in CD11c+ cells in whole body metabolism regulation by controlling norepinephrine levels in WAT, modulating catecholamine-induced lipolysis and increasing thermogenesis that together created a lean phenotype.

## INTRODUCTION

Obesity is associated with a state of chronic, low-grade inflammation that contributes to insulin resistance and susceptibility to type II diabetes in rodents and humans^1^ and recent studies have revealed that the immune system and metabolism are highly integrated. Diet-induced obesity is characterized by the recruitment of immune cells, including CD11c+ macrophages and dendritic cells, into adipose tissue^2^. CD11c (*Itgax*) is a type I transmembrane protein found on hematopoietic cells, such as on most dendritic cells (DCs), monocytes, macrophages, neutrophils, B and T cells as well as NK cells, in rodents and humans^3–5^. Tissue resident immune cells in a lean animal generally have an anti-inflammatory phenotype and diet-induced obesity (DIO) induces trafficking of CD11c+ monocyte-derived macrophages and DC into adipose tissue^6^. These recruited CD11c+ immune cells are important factors in the pathogenesis of metabolic disease since their ablation attenuates adipose tissue inflammation and improves glucose tolerance without changes in body weight^7^.

The G-protein Gas (encoded by *Gnas*) regulates production of cyclic AMP (cAMP) and mediates the sympathetic nervous system (SNS) effects on brown adipose tissue (BAT) to stimulate thermogenesis and on white adipose tissue (WAT) to promote lipolysis^8^. Denervation of adipose tissue and reduces lipolysis and thermogenesis^9, 10^ underscoring the importance of the SNS in regulating obesity and energy expenditure. The catecholamine norepinephrine (NE) is the major effector of the SNS and it stimulates adaptive thermogenesis and lipolysis through b-adrenergic receptors^11^. Mice lacking all b-adrenergic receptors (b-ARs) develop massive obesity due to a defect in stimulated lipolysis and a deficiency in diet-induced thermogenesis^12^. Activation of BAT to increase energy expenditure via uncoupled thermogenesis is an attractive anti-obesity strategy^13^. Recently, brown adipocyte-like cells, also referred to as beige or brite cells, have been described within WAT under thermogenic conditions^14^. Such beige adipocytes display similar physiological functions of brown adipocytes but are derived from a different lineage. Beige adipocytes can also express uncoupling protein 1 (UCP1) to allow thermogenesis. Given the higher mass of WAT compared to BAT, particularly in humans, beigeing of white adipose tissue could have anti-obesity effects.

Here, we demonstrate that disruption of Gas signaling in CD11c+ immune cells improved glucose metabolism in mice by promoting energy expenditure and inducing adipose beigeing, and consequently prevented obesity and insulin resistance. Notably, we identified an important role for Gas in CD11c+ immune cells to regulate adipose tissue catecholamine levels that likely contributed to the lean phenotype that constitutes a potential target for the treatment of obesity and T2D.

## RESULTS

### *Gnas^ΔCD11c^* KO mice are resistant to diet-induced obesity and insulin resistance

*Gnas^ΔCD11c^* mice were generated as previously described^15^. We first monitored the body weights (BW) of *Gnas^ΔCD11c^* (KO) and *Gnas^fl/fl^* (Flox) mice on a normal chow diet (NCD) and found that both male and female KO mice exhibited significantly decreased body weight when compared with control littermate Flox mice **(Fig. 1a)**. Tissue weights were similar except that eWAT was slightly lighter in the KO mice (**Supplemental Fig. 1a**). Liver, brown adipose tissue (BAT), epididymal or inguinal white adipose tissue (eWAT or iWAT) histologies were indistinguishable between genotypes, and adipocyte size distribution was the same (**Supplemental Fig. 1b and c**). At 5 months of age, KO mice showed improved glucose tolerance **(Fig. 1b)** and insulin sensitivity **(Fig. 1c**), and lower fasting insulin **(Fig. 1d)** and glucose levels **(Fig. 1e)**, and consequently lower HOMA-IR (homeostatic model of insulin resistance) **(Fig. 1f)** compared to controls. To further address whether Gas deficiency in CD11c+ cells influenced energy homeostasis, the mice were placed in metabolic cages. The body weight-matched KO mice manifested comparable food intake, oxygen consumption rate (VO_2_) and energy expenditure (EE) as control mice under NCD **(Fig. 1g-i).** No significant difference in VCO_2_ or respiratory exchange ratio (RER) were observed but the KO mice had slightly higher physical activity and drinking in the dark phase **(Supplemental Fig. 1d-h)**.

**Figure 1:**
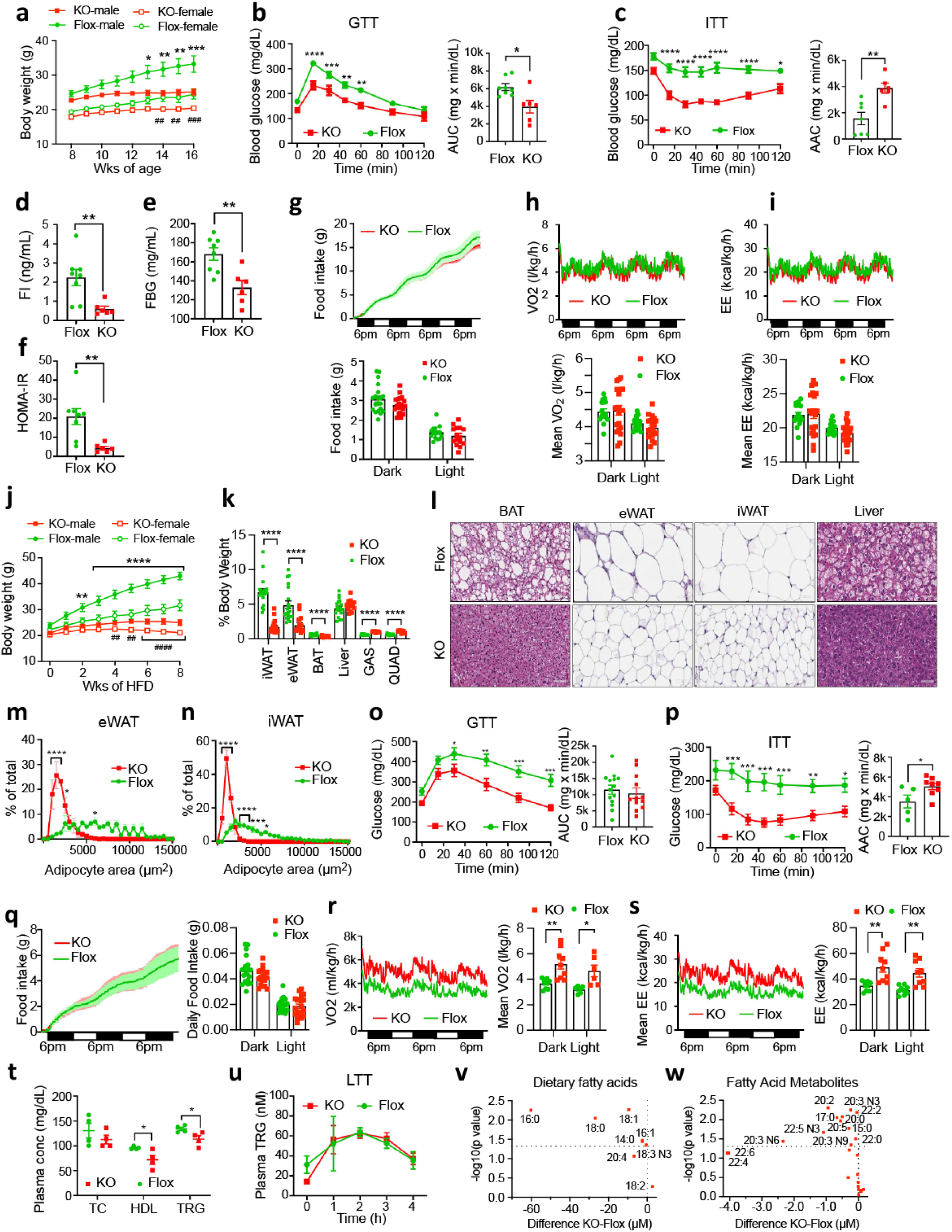
Gnas^ΔCD11c^ mice are protected from diet-induced obesity (DIO) For all panels *Gnas^fl/fl^* (Flox) mice are shown in green circles, *Gnas^DCD11c^* (KO) mice in red squares. (**a**) Body weights of male and female littermates on normal chow diet (NCD) from age 8 to 16 weeks (n=8-10/group). (**b**) Glucose tolerance tests of mice on NCD. Bar graph show area under the curve (AUC). (**c**) Insulin tolerance tests of mice on NCD. Bar graph shows area above the curve (AAC). (**d, e, f**) Fasting insulin and glucose levels and HOMA-IR of Flox and KO on NCD for 20 weeks. (**g**-**i**) Metabolic assessment of Flox and KO mice on NCD; (**g**) cumulative food intake (top) and average food intake during dark or light cycle (bottom); (**h**) oxygen consumption (VO_2_) over time (top), and average VO_2_ during light and dark cycle (bottom); (**i**) energy expenditure (EE) over time (top) and average EE during dark and light cycle (bottom). (**j**) Body weights of male and female, Flox and KO mice on 60% high-fat diet (HFD) were recorded for 8 weeks starting at 8 weeks of age (n=19-20 for males and n=8-12 for females). Asterisks indicate statistically significant genotype differences for male mice, hashtags indicate significant differences for female mice. (**k**) Tissue weights (expressed as percent body weight) of Flox and KO mice after 8 weeks of HFD feeding. iWAT, eWAT, brown adipose tissue (BAT), liver, gastrocnemius muscle (GAS), and quadriceps muscle (QUAD). (**l**) Representative H&E staining of BAT, eWAT, iWAT and liver of HFD-fed Flox and KO mice. Scale bar=100 mm. (**m** and **n**) Adipocyte area distribution in eWAT (left) and iWAT (right) from male Flox and KO mice on HFD. (**o**) GTT for mice on HFD and area under the curve (AUC). Time (p<0.0001), genotype (p=0.0029), and interaction (p=0.055) by repeated measures 2-way ANOVA. (**p**) ITT for mice on HFD and area above the curve (AAC). (**q**-**s**) Metabolic assessment of HFD-fed Flox and KO mice; (**q**) cumulative food intake (left) and daily food intake during dark and light phases (right); (**r**) oxygen consumption rate (VO_2_) over time (left), average VO_2_ during dark and light phases (right); (**s**) energy expenditure (EE) over time (left) and average EE during dark and light phases (right). (**t**) Plasma total cholesterol (TC), high-density lipoprotein cholesterol (HDL-C) and triglyceride (TRG) levels of HFD-fed Flox and KO mice were determined using Cholestech LDX system. n=4/group. (**u**) Oral lipid tolerance test (Oral LTT), triglyceride content of Flox and KO mice fasted overnight in response to oral fat load. n=4/group. (**v**) Volcano plot of plasma dietary fatty acids for Flox and KO mice on HFD. Plot shows the difference in FA concentration (µM) against the -log10 p-value. (**w**) Volcano plot of plasma fatty acid metabolites for Flox and KO on HFD (n=4/group). Data are represented as mean ± SEM and asterisks indicate Statistical significance by 2-way repeated measures ANOVA or unpaired Student’s t-test as appropriate; *p < 0.05; **p < 0.01; ***p < 0.001.

We next challenged the mice with 60% kcal high fat diet (HFD). Both male and female KO mice showed resistance to HFD-induced body weight gain **(Fig. 1j)**. The iWAT, eWAT, BAT and liver were significantly lighter in the KO mice, but the spleen was slightly heavier **(Fig. 1k and Supplemental Fig. 1i)**. HFD caused the expected whitening of the BAT, adipocyte enlargement in eWAT and iWAT, and hepatic steatosis in the Flox mice, but the KO mice did not show any of these changes (**Fig. 1l-n**). Consistent with their lean phenotype, KO mice showed improved glucose tolerance **(Fig. 1o)** and insulin sensitivity **(Fig. 1p)**. The mice on HFD were also subjected to a metabolic analysis. Neither cumulative nor daily food and water intake was different between the groups (**Fig. 1q and Supplemental Fig. 1j**) but BW-adjusted VO_2_, EE and VCO_2_ were much higher in the KO mice during both the light and dark phases (**Fig. 1r and s, and Supplemental Fig. 1k**) and showed a differential dependence on body weight (**Supplemental Fig. 1l**). These changes were not associated with increased RER or physical activity **(Supplemental Fig. 1m-o)**. Next, we evaluated the lipid metabolism in the mice and found KO mice had lower levels of high-density lipoprotein cholesterol (HDL) and triglycerides (TRG) in the serum as compared with their littermate controls **(Fig. 1t)**. To determine if this difference in body weight is associated with deficiency in lipid absorption, we performed a lipid tolerance test and gave the mice a bolus of sesame oil by oral gavage. The KO mice had comparable lipid tolerance **(Fig. 1u)**. As serum lipid levels were reduced in the KO mice, we compared the fatty acid (FA) profiles. The KO mice had lower levels of palmitic (16:0), stearic (18:0) and oleic (18:1) acids (**Fig. 1v**) which are the major components of the HFD and are increased in obese mice (**Supplemental Fig. 1p**), as well as lower polyunsaturated FA metabolites including eicosadienoic (20:2), di-homo-g-linoleic (20:3N6), docosapentaenoic (22:5N3), adrenic (22:4), and docosahexaenoic (22:6) acid (**Fig. 1w**) including some that are increased in obesity (**Supplemental Fig. 1q**). These results collectively demonstrate that the *Gnas* ablation in CD11c+ cells protected mice from obesity and from impaired glucose and insulin tolerance.

The dramatic increase in VO_2_ observed in the HFD-fed KO mice led us to the question that whether the difference in VO_2_ is triggered by diet-induced thermogenesis with the HFD. To address this question, we fed body weight-matched KO and Flox mice with normal chow diet for 3 days and then switched to HFD for another 3 days while monitoring whole body metabolism by CLAMS (**Supplemental Fig. 2a**). As expected, the RER for mice on NCD showed circadian rhythms with alternating utilization of carbohydrate during the dark phase and lipid during the light phase (**Supplemental Fig. 2b**), and after the switch to HFD the RER decreased, and the circadian changes were suppressed. There were no differences in RER with genotype, but the RER was lower on HFD during the dark phase (**Supplemental Fig. 2c**) but not the light phase (**Supplemental Fig. 2d**). There was no genotype difference in food intake (**Supplemental Fig. 2e-g**) or water consumption (**Supplemental Fig. 2h-j**) either before or after the switch to HFD. VO_2_ was increased by HFD in both genotypes in both the light and dark phases (**Supplemental Fig. 2k-m**) as was VCO_2_ (**Supplemental Fig. 2n-p**). No differences were observed in physical activity **(Supplementary Fig. 2q-v)**. Taken together, the results suggest that the difference in VO_2_ was not triggered by diet-induced thermogenesis.

### Immune cells are responsible for the lean phenotype of *Gnas^ΔCD11c^* KO mice

CD11c is expressed on mouse immune cells including dendritic cells (DCs), monocytes/macrophages, neutrophils and natural killer (NK) cells, but is also expressed on microglia in the brain^6^. To assess the role of hematopoietic cells in the lean phenotype, we generated chimeric mice by transplanting bone marrow from KO donor mice into lethally irradiated obese wild-type (WT) C57BL/6J recipient mice. As a control, we transplanted bone marrow from Flox mice into irradiated obese C57BL/6J recipient mice **(Fig. 2a)**. We verified the chimerism 6 weeks after bone marrow transplantation (BMT) by q-PCR and PCR analysis for the relative levels of *Cre* and *Gnas^fl/fl^* in genomic DNA of peripheral blood cells. We observed that mice receiving the KO bone marrow start to lose body weight 7 weeks after transplantation once the immune system has been reconstituted compared to mice receiving WT bone marrow **(Fig. 2b)** despite no change in food intake **(Fig. 2c)**. The Flox chimeric mice did not show improved glucose tolerance at 7 weeks (**Fig. 2d**), but the KO chimeric mice displayed improved glucose tolerance and insulin sensitivity at weeks 8 and 9 after BMT **(Fig. 2e and f)** which was still evident at week 11 (**Supplemental Fig. 3a**). These results suggested that lack of Gas in CD11c+ immune cells led to metabolic improvement and resistance to DIO.

**Figure 2:**
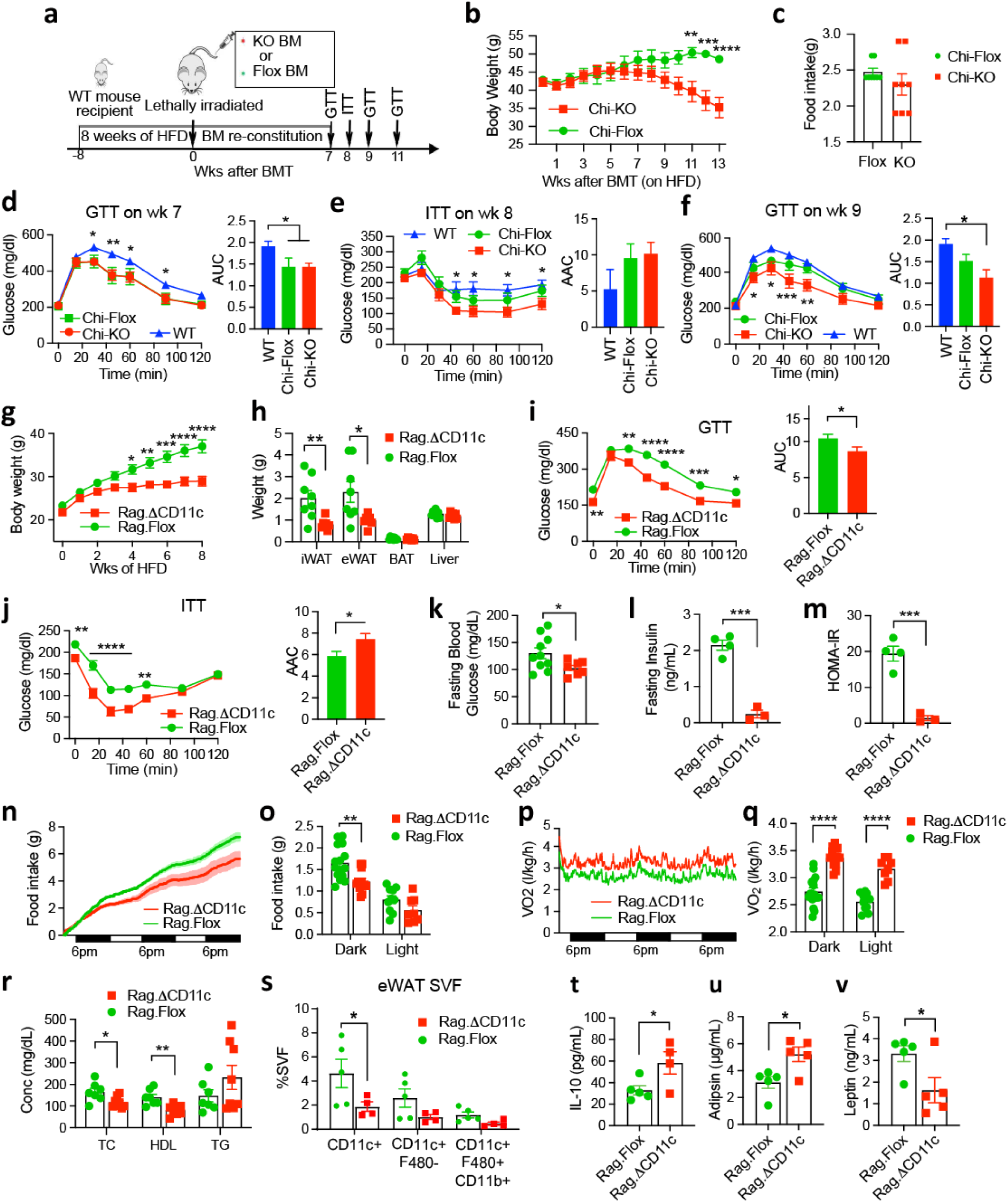
Immune cells are responsible for the lean phenotype of Gnas^ΔCD11c^ KO mice. (a) Schematic of bone marrow transplantation experiment into obese recipient mice on HFD. Body weight of chimeric mice after engraftment, mice received bone marrow cells from KO mice (Chi-KO) or Flox mice (Chi-Flox). (**c**) Daily food intake for Chi-KO and Chi-Flox recipients. (**d** and **f**) Glucose tolerance tests performed 7 weeks and 9 weeks after transplantation for Chi-KO and Chi-Flox mice compared to control obese WT mice (n=8/group). Area under the curve is shown on right. (**e**) Insulin tolerance test for Chi-KO and Chi-Flox mice compared to obese WT mice (n=8/group). Area above the curve is shown on right. (**g**) Body weights of *Rag1:Gnas^DCD11c^* (Rag.DCD11c) and *Rag1:Gnas^fl/fl^* (Rag.Flox) mice fed a 60% HFD (n=11-15/group). (**h**) Tissue weights for iWAT, eWAT, BAT and liver of Rag.DCD11c and Rag.Flox mice after 8 weeks of HFD feeding (n=5-8/group). (**i** and **j**) GTTs and ITTs on Rag.DCD11c and Rag.Flox mice after 8 weeks of HFD (n=6-15/group). AUC and AAC are shown in the right-hand panels. Fasting glucose (**k**), fasting insulin (**l**) and HOMA-IR (**m**) from HFD-fed Rag.Flox and Rag.DCD11c mice. (**n**) Cumulative food intake and (**o**) average daily food intake during dark and light cycle during CLAMS analysis. (**p**) Oxygen consumption rate (VO_2_) over time and (**q**) average VO_2_ during light and dark cycle. For CLAMS analysis n=5 for Rag.Flox and n=4 for Rag.DCD11c. (**r**) Blood TC, HDL and TRG levels of HFD-fed Rag.Flox and Rag.DCD11c mice. (**s**) Stromo-vascular cells (SVCs) were extracted from the eWAT samples and analyzed by FACS. The number of CD11c+, CD11c+F480- and F4/80+CD11b+CD11c+ cells is indicated as percentage of total SVCs (n=5 for Rag.Flox group and n=4 for Rag.DCD11c group). (**t**-**v**) Serum levels of IL10 (**t**), adipsin (**u**) and leptin (**v**) in HFD-fed Rag.Flox and Rag.DCD11c mice (n=5/group). For all graphs, data are presented as mean ± SEM and asterisks indicate statistical significance by unpaired Student’s t-test or repeated measures 2-way ANOVA as appropriate; *p<0.05, **p<0.01, ***p<0.001 and ****p<0.0001.

To determine whether the ablation of *Gnas* in CD11c cells plays a role in the metabolic improvement in mice by initiating adaptive T cell responses, we generated *Rag1^-/-^/Gnas^ΔCD11c^* double KO mice (Rag.ΔCD11c) by crossbreeding the *Gnas^ΔCD11c^* mice with *Rag1*-deficient mice that lack T and B lymphocytes. The Rag.ΔCD11c mice were resistant to DIO when fed HFD (**Fig. 2g**) and consequently had reduced iWAT and eWAT (**Fig. 2h**). The Rag.DCD11c mice showed decreased lipid accumulation in BAT, and smaller adipocyte size in eWAT and iWAT (**Supplemental Figure 3b**). The Rag.ΔCD11c mice also displayed improved glucose tolerance and insulin sensitivity (**Fig. 2i and j**) and had decreased fasting blood glucose (**Fig. 2k**), insulin levels (**Fig. 2l**), and HOMA-IR (**Fig. 2m**) relative to Rag.Flox controls. We measured whole-body metabolism by CLAMS. The Rag.DCD11c mice were lighter than the Rag.Floxcontrols before and after the study (**Supplemental Fig. 3c**) and had reduced food intake (**Fig. 2n and o**) but increased BW-adjusted VO_2_ **(Fig. 2p** and **q**), EE and VCO_2_ **(Supplemental Fig. 3d and e**), independent of BW (**Supplemental Fig. 3f**). No significant differences were observed in terms of drinking, RER and ambulatory activity between the two groups (**Supplemental Fig. 3g-j**), suggesting that elevated energy expenditure and decreased appetite lead to less body weight gain in the Rag.ΔCD11c mice.

Diet-induced obesity leads to an increase in M1-like adipose tissue macrophages (ATMs) displaying the CD11c surface marker^2^. To directly assess the participation of macrophages in the KO metabolic phenotype, we deleted *Gnas* in macrophages using LysM-Cre. Male *Gnas^ΔLysM^* mice (LysM-KO) and control littermates (Flox) were placed on a 60% high fat diet at 8 weeks of age for 13 weeks. LysM-KO mice gained comparable body weight to Flox control mice (**Supplemental Fig. 4a**). Glucose tolerance and insulin sensitivity were not significantly different between the groups (**Supplemental Fig. 4b and c**). The results suggested that the CD11c+ve cells responsible for the lean phenotype do not express the LysM-Cre driver^16^.

Numerous studies have demonstrated that the gut microbiota plays an important role in the induction of obesity^17^. To examine the involvement of the microbiota in the resistance to obesity in *Gnas^ΔCD11c^* mice, we co-housed the KO mice together with their littermate Flox mice and found the KO mice maintained the lean phenotype on HFD feeding while the Flox mice became obese (**Supplemental Fig. 4d**). To further test the role of microbiota in the KO mice, we transplanted feces from KO and Flox mice to germ-free WT mice (**Supplemental Fig. 4e**). The mice that received feces from KO mice displayed comparable body weight gain with the mice receiving that from Flox mice (**Supplemental Fig. 4f**). These data collectively indicate that the lean phenotype in the KO mice cannot be attributed to alterations in the gut microbiota.

### *Gnas^ΔCD11c^* KO mice have reduced adipose tissue inflammation

Obesity is associated with chronic inflammation characterized by progressive accumulation of immune cells in adipose tissue^2^. We therefore quantified the expression of immune cells marker genes and several genes related to inflammation in epididymal adipose tissue (eWAT) by qPCR. As expected, HFD led to increase in the expression of the macrophage markers *Itgax* (CD11c) and *Adgre1* (F4/80), as well as the inflammatory cytokines *Il6* and *Ccl2* (MCP1), and this increase was markedly blunted in the KO mice (**Fig. 3a**). Interestingly, the macrophage marker *Itgam* (CD11b) and *Il10* were higher in the KO and further increased in response to HFD. We also measured circulating cytokines. IL-6 was increased in the KO mice on HFD and, notably, the anti-inflammatory cytokine IL-10 was 50-fold higher in the KO mice as compared to Flox mice (**Fig. 3b**). No significant differences in the levels of TNFa, IL1b and CCL2 were observed between the groups (**Supplemental Fig. 5a-e**). We also observed fewer crown-like structure in the eWATs of KO mice (**Fig. 3c and d**) consistent with decreased adipose inflammation. We further analyzed the expression of the adipokines adipsin, adiponectin and leptin. Expression of adipsin and adiponectin in eWAT by qPCR decreased after HFD feeding in controls, while KO mice had slightly lower levels and did not respond to HFD (**Fig. 3e and Supplemental Fig. 5f**), and expression of leptin in eWAT was lower in the KO (**Fig. 3f**). Serum adipsin levels were significantly decreased in HFD-fed animals in both genotypes, and it was significantly higher in KO on HFD than the Flox-HFD control group (**Fig. 3g**). Conversely, the serum leptin levels were dramatically elevated in obese Flox mice, while it was not elevated in the KO mice (**Fig. 3h**), which is consistent with the lean phenotype. Serum adiponectin was decrease by HFD in the KO but brain-derived neutotrophic factor (BDNF) was not changed (**Supplemental Fig. 5g and h**). We also assessed adipose tissue from the Rag.DCD11c mice. The numbers of adipose tissue CD11c+ cells, CD11c+/F4/80- cells, and CD11c+/F4/80+/CD11b+ macrophages were unchanged in the Rag.DCD11c mice (**Supplemental Fig. 6a**). There were no differences in serum cytokines, adiponectin or BDNF (**Supplemental Fig. 6b and c**).

**Figure 3:**
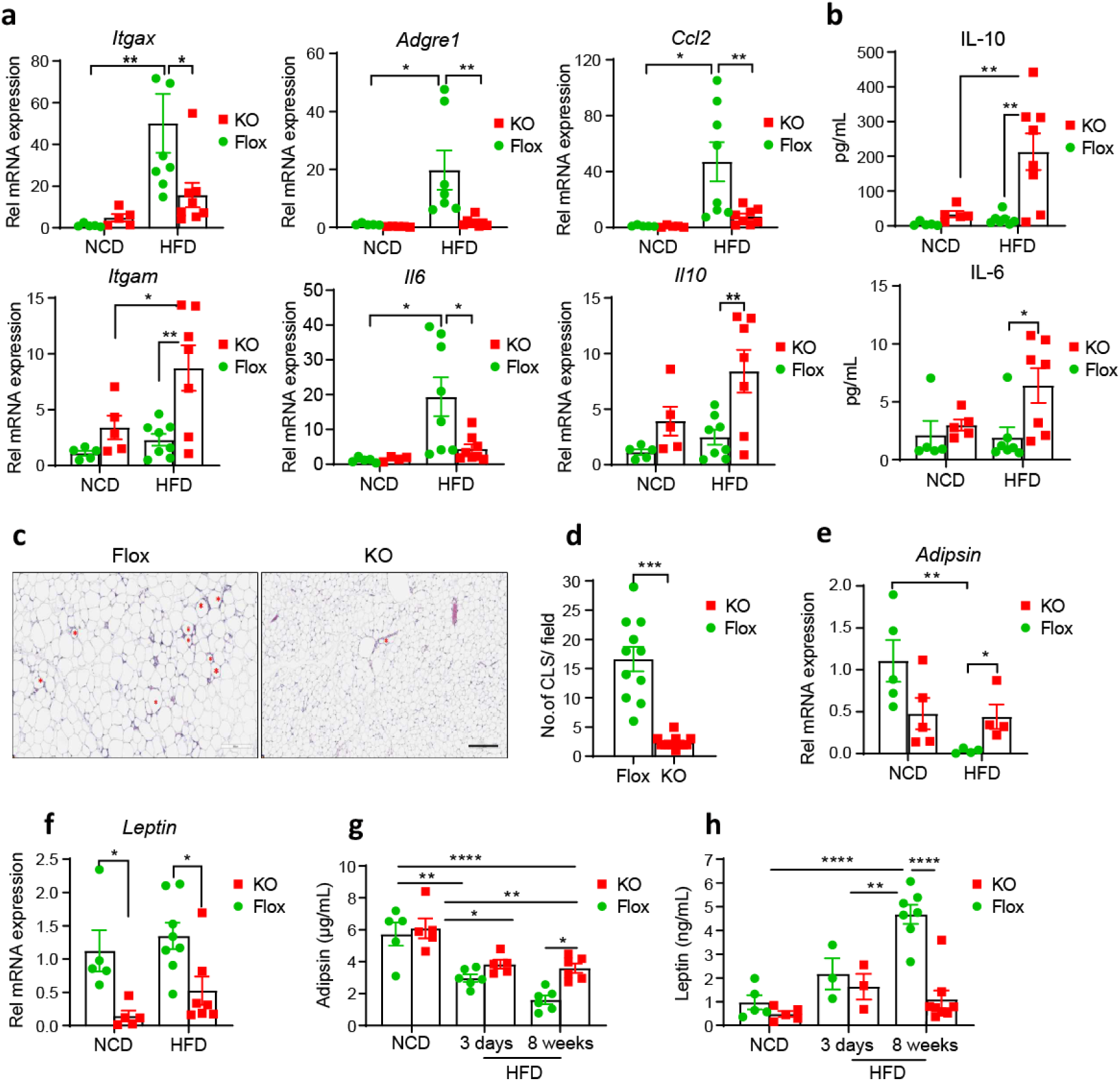
Gnas^ΔCD11c^ KO mice have reduced adipose tissue inflammation. (**a**) Gene expression of *Itgax* (CD11c)*, Itgam* (CD11b), *Adgre1* (F4/80), *Il6* (interleukin-6), and *Ccl2* (MCP-1), and *Il10* (interleukin-10) in eWAT from KO and Flox mice fed with NCD or HFD for 8 weeks by qPCR. Asterisks indicate significance by 2-way ANOVA (n=5-8/group). (**b**) Plasma protein levels of IL6 and IL-10 for Flox and KO mice (n=5-8/group). (**c**) Representative H&E staining of sections of eWAT tissue. Crown-like structures (CLS) are indicated with red asterisks. (**d**) Quantification of CLS density of the H&E-stained sections (n=4/group). (**e and f**) Relative mRNA levels of adipsin (**e**) and leptin (**f**) in eWATs of HFD-fed Flox and KO mice fed with NCD or HFD as indicated, 36B4 was used as loading control (n=5-8/group). (**g and h**) Adipsin (**g**) and leptin (**h**) protein levels in the serum of Flox and KO mice fed with NCD or HFD as indicated were determined by ELISA (n=5-7/group). Data are presented as means ± SEM and asterisks indicate statistical significance by unpaired Student’s t-test or 2-way ANOVA as appropriate; *p<0.05, **p<0.01, ***p<0.001, ****p<0.0001.

### Energy expenditure is increased in *Gnas^ΔCD11c^* KO mice

As energy expenditure is increased in KO mice on HFD, we investigated the thermogenic activity of Flox and KO mice on HFD using infrared imaging of interscapular BAT. We observed increased dorsal interscapular BAT temperature in the KO mice as compared with Flox mice (**Fig. 4a**). No significant changes were found between the NCD groups but the BAT in KO mice on HFD was almost 2°C warmer **(Fig. 4b)**. To further investigate the phenotype, we measured oxygen consumption in several tissues in response to specific substrates and inhibitors using an Oroboros-2k system. No differences were observed in livers of mice on NCD (**Fig. 4c**), but combined complex I and II activity was higher in livers of KO mice on HFD (**Fig. 4d**). Complex I activity was higher in soleus muscles of KO mice on NCD (**Fig. 4e**), while complex I and II activity and maximal respiration were increased in soleus muscles from KO mice on HFD (**Fig 4f**). No alterations in oxygen consumption in WAT, BAT and gastrocnemius skeletal muscle were observed; however the values were at the limit of detection (**Supplemental Fig. 7a-h**). We then addressed whether the KO mice could increase thermogenesis in response to a cold challenge. Mice were placed in a cold room at 4°C and body temperature measured using subcutaneous temperature sensors. The Flox mice were able to maintain their body temperature over > 4 h, but the body temperature of KO mice started to drop after 3 h of cold exposure and had dropped by almost 5°C after 4.5 h. Body temperature quickly recovered when mice were removed from the cold room (**Fig. 4g**). This suggested that the KO mice were unable to respond to the sympathetic activation of thermogenesis. To test whether the immune cells could be contributing to the thermogenesis in these mice, we measured oxygen consumption in splenic cDC2 cells and naïve CD4 T cells by Seahorse XF. The cDC2 cells from the KO mice exhibited marked higher basal oxygen consumption rate (OCR) along with increased maximal OCR (**Fig. 4h** and **i**). In contrast, the CD4 T cells showed comparable basal and maximal OCR between the KO and Flox controls (**Fig. 4j** and **k**). So, the increased oxygen consumption in CD11c cells could contribute to the lean phenotype.

**Figure 4:**
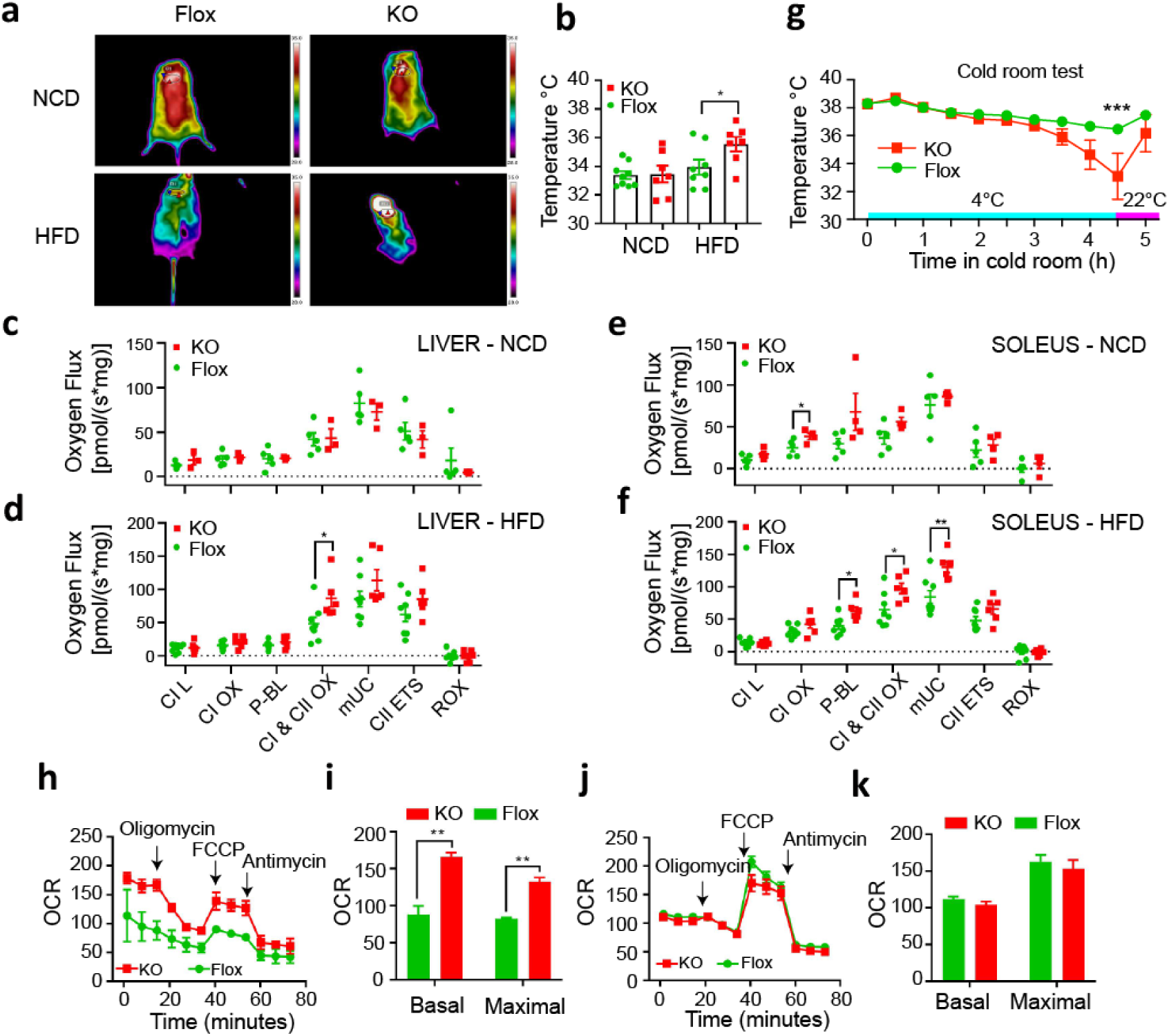
Energy expenditure is increased in Gnas^ΔCD11c^ KO mice. Infrared thermography was used to quantify heat generation from the interscapular brown fat depot. (**a**) Thermal camera images of dorsal view of Flox and KO mice fed with indicated diet (NCD or HFD). (**b**) Quantification of BAT temperatures of mice from representative images (n=7-9 mice/group). Oroboros tissue O_2_ fluxes in liver (**c** and **d**) and soleus muscle (**e** and **f**) of Flox and KO mice on indicated diet with complex 1 substrates glutamate and malate (CI L), with ADP added to allow oxidative phosphorylation (CI Ox), followed by addition of pyruvate (P-BL), then CII substrate succinate (CI & CII Ox), then FCCP to uncouple mitochondria (mUC), then addition of rotenone to inhibit CI to assess CII uncoupled (CII ETS), and malonate/antimycin to inhibit CII and CIII activity to assess residual oxygen consumption (ROX). n=3-5 for NCD groups, n=6-8 for HFD groups. (**g**) Body temperatures of NCD-fed Flox and KO mice with subcutaneous temperature sensors placed in cold room at 4°C (n=5/group). Oxygen-consumption rate (OCR) of FACS-sorted splenic cDC2 (**h** and **i**) and splenic CD4+ T cells (**j** and **k**) of HFD-fed KO and Flox mice by Seahorse analysis under basal conditions followed by the sequential addition of oligomycin, FCCP, and antimycin/rotenone (n=2-3/group). Data are presented as means ± SEM and asterisks indicate statistical significance by unpaired Student’s t-test, or 1-way or 2-way ANOVA as appropriate; *p<0.05, **p<0.01, ***p<0.001.

### Loss of *Gnas* in CD11c+ cells increases cAMP signaling and thermogenic capacity of adipose tissue

Mitochondrial uncoupling protein 1 (UCP1) is responsible for non-shivering thermogenesis in brown adipose tissue by increasing the conductance of the inner mitochondrial membrane to dissipate membrane potential so that the mitochondria generate heat rather than ATP. As thermogenesis was increased in the KO mice, we measured UCP1 protein in BAT, eWAT and iWAT of KO mice (**Fig. 5a**). UCP1 protein was reduced in BAT but increased in eWAT and iWAT (**Fig. 5b**) suggesting that the increased energy expenditure in the KO mice is most likely the result of beigeing of WAT. So, we measured the expression of brown and beige associated genes in BAT, eWAT and iWAT in these mice. *Ucp1* mRNA was reduced in BAT consistent with the protein data (**Fig. 5c**), as was expression of *Oplah, Pdk4* and *Serca2b*. Expression of *Ucp1* was highly elevated in eWAT, as was *Pgc1a, Cidea* and *Pparg* (**Fig. 5d**). *Cidea* and *Pparg* were elevated in iWAT and *Ucp1* tended to increase but was not significant (**Fig. 5e**). Similar changes were seen in BAT, eWAT and iWAT from the obese Rag.DCD11c mice (**Supplemental Fig. 8a-d**). The thermogenic pathway is activated by cAMP signaling, so we measured activation of PKA in eWAT and iWAT by immunoblotting with an antibody to phospho-PKA substrates. Several proteins showed increased phosphorylation in eWAT and iWAT (**Fig. 5f-i**), so we then measured the phosphorylation of hormone sensitive lipase (HSL/LIPE) in the eWAT and iWAT of KO and Flox mice on HFD. Phosphorylation of HSL on Ser563 was increased in both iWATs and eWAT (**Fig. 5j-k**). HSL is the major lipase involved in stimulated lipolysis, whereas adipose triglyceride lipase (ATGL/PNPLA2) is the major lipase involved in basal lipolysis^18^. Both eWAT and iWAT showed high expression of ATGL protein in the KO mice on HFD (**Fig. 5j-k**). Similar changes were observed in eWAT and iWAT from the obese Rag.DCD11c mice (**Supplemental Fig. 8e**). cAMP signaling can inhibit the activity of mammalian target of rapamycin complex 1 (mTORC1) so we measured phosphorylation of mTOR targets. As expected, we detected a significant decrease in phosphorylation of S6 kinase (p-S6K) in the iWAT of KO mice, in relative to Flox mice (**Fig. 5l and m**). Similar reductions in mTOR activity were observed in the obese Rag.DCD11c mice (**Supplemental Fig. 8f)**. We did not see phosphorylation of CREB, HSL or induction of UCP1 or ATGL in either lean KO or Flox mice on NCD (**Supplemental Fig. 9a**).

**Figure 5:**
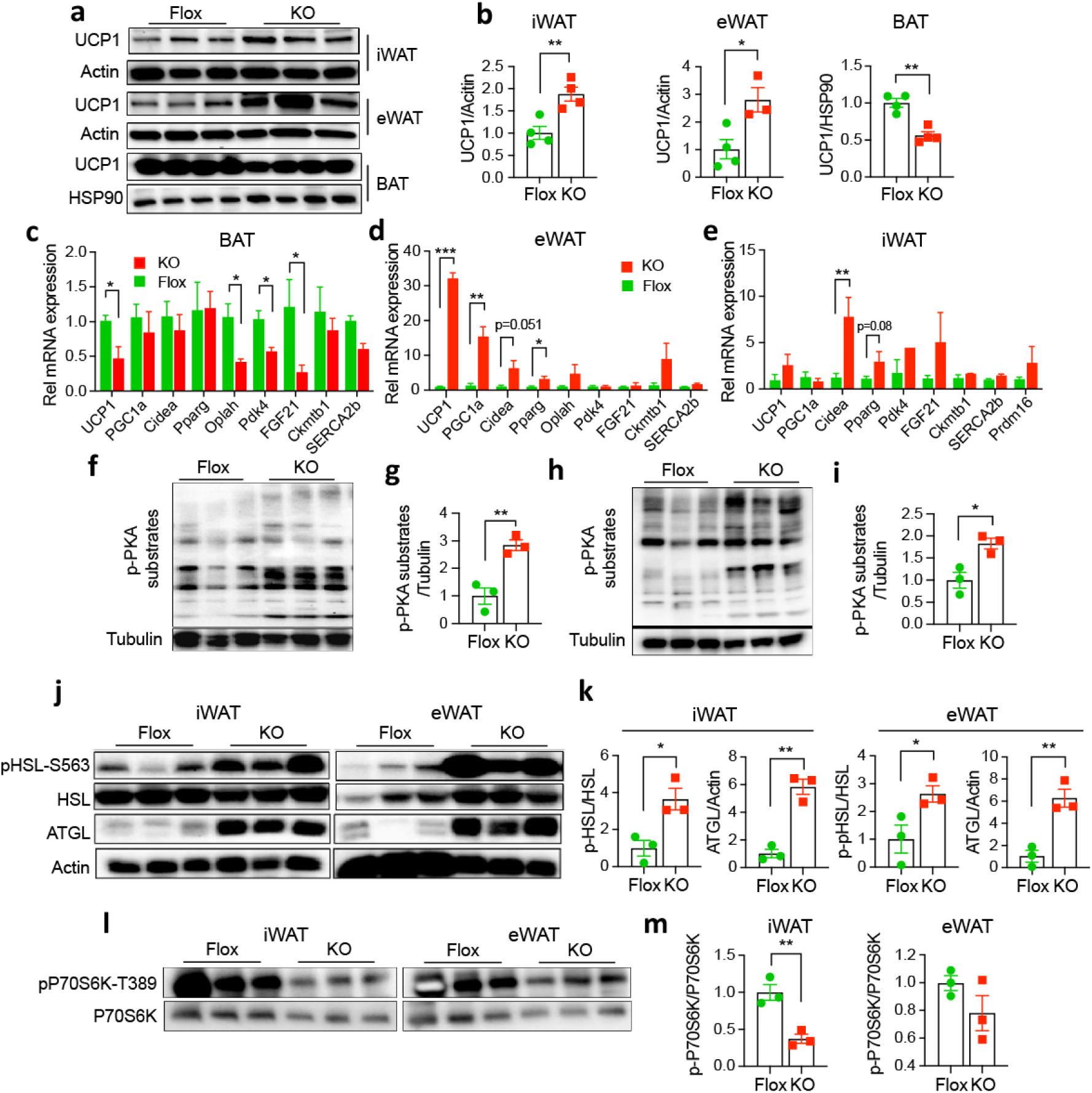
Loss of Gnas in CD11c+ cells increases cAMP signaling and thermogenic capacity of adipose tissue. (**a**) Representative immunoblots of UCP1 in iWAT, eWAT and BAT of HFD-fed Flox and KO mice. Actin and HSP90 were used as loading controls. (**b**) Quantification of UCP1 western data. (**c**-**e**) Relative expression of beige-related genes in BAT (**c**), eWAT (**d**) and iWAT (**e**) of HFD-fed Flox and KO mice normalized to m36B4 control mRNA (n=3-4/group). (**f** and **h**) Representative immunoblots of PKA kinase activity using an anti-phospho-PKA substrate antibody in iWAT (**f**) and eWAT (**h**) extracts from HFD-fed Flox and KO mice. Tubulin was used as a loading control. (**g** and **i**) Quantification of p-PKA substrates normalized to Tubulin in iWAT and eWAT. (**j**) Immunoblots for phospho-Ser563 HSL, total HSL, ATGL and tubulin in iWAT and eWAT extracts from HFD-fed Flox and KO mice. (**k**) Quantification of pHSL-S563/HSL and ATGL/Tubulin in iWAT and eWAT (n=3-4/group). (**l**) Immunoblots phosphor-Thr389 p70S6K, total p70S6K in iWAT and eWAT extracts from HFD-fed Flox and KO mice. (**m**) Quantification of p-P70S6K-T389 to total P70S6K in iWAT and eWAT (n=3-4/group). Data are presented as means ± SEM and asterisks indicate statistical significance by unpaired Student’s t-test or ANOVA as appropriate; *p<0.05, **p<0.01.

### Norepinephrine and dopamine are elevated in WAT of GNAS^ΔCD11c^ KO mice

Based on the observation of increased cAMP signaling in adipose tissue, we reasoned that deficiency of *Gnas* in CD11c cells could cause upregulation of Gas-linked GPCR ligands, which could activate PKA signaling in neighboring adipocytes. To test this hypothesis, we treated HEK293 cells with serum from either KO or Flox mice and found that the KO serum could induce greater phosphorylation of PKA substrates and the cAMP-dependent transcription factors CREB and ATF1 compared to Flox serum (**Fig. 6a** and **b**). We also detected higher intra-cellular cAMP in the cells treated with KO serum, relative to Flox serum, and the increase of cAMP was blunted when the cells were pretreated with propranolol (**Fig. 6c**), suggesting that the effect was mediated by the activation of b-adrenergic receptors (bARs). There are three Gas-coupled bAR subtypes (b1AR, b2AR and b3AR). b1AR and b2AR are broadly expressed throughout the body, while b3AR is found predominantly in adipocytes and is involved in the regulation of lipolysis and thermogenesis. So, we compared the expression levels of bARs in the WAT of KO and Flox mice under two different diets, NCD and HFD. As expected, b3AR transcripts (*Adrb3*) were decreased dramatically in both eWAT and iWAT of Flox mice when fed the HFD, as compared to NCD. The expression of b3AR was partially reduced in the KO WAT on NCD and did not change upon HFD-feeding (**Fig. 6d**). In contrast, there were no changes in the expression levels of b1AR (*Adrb1*) and b2AR (*Adrb2*) between the groups (**Supplemental Fig. 10a and b**).

**Figure 6:**
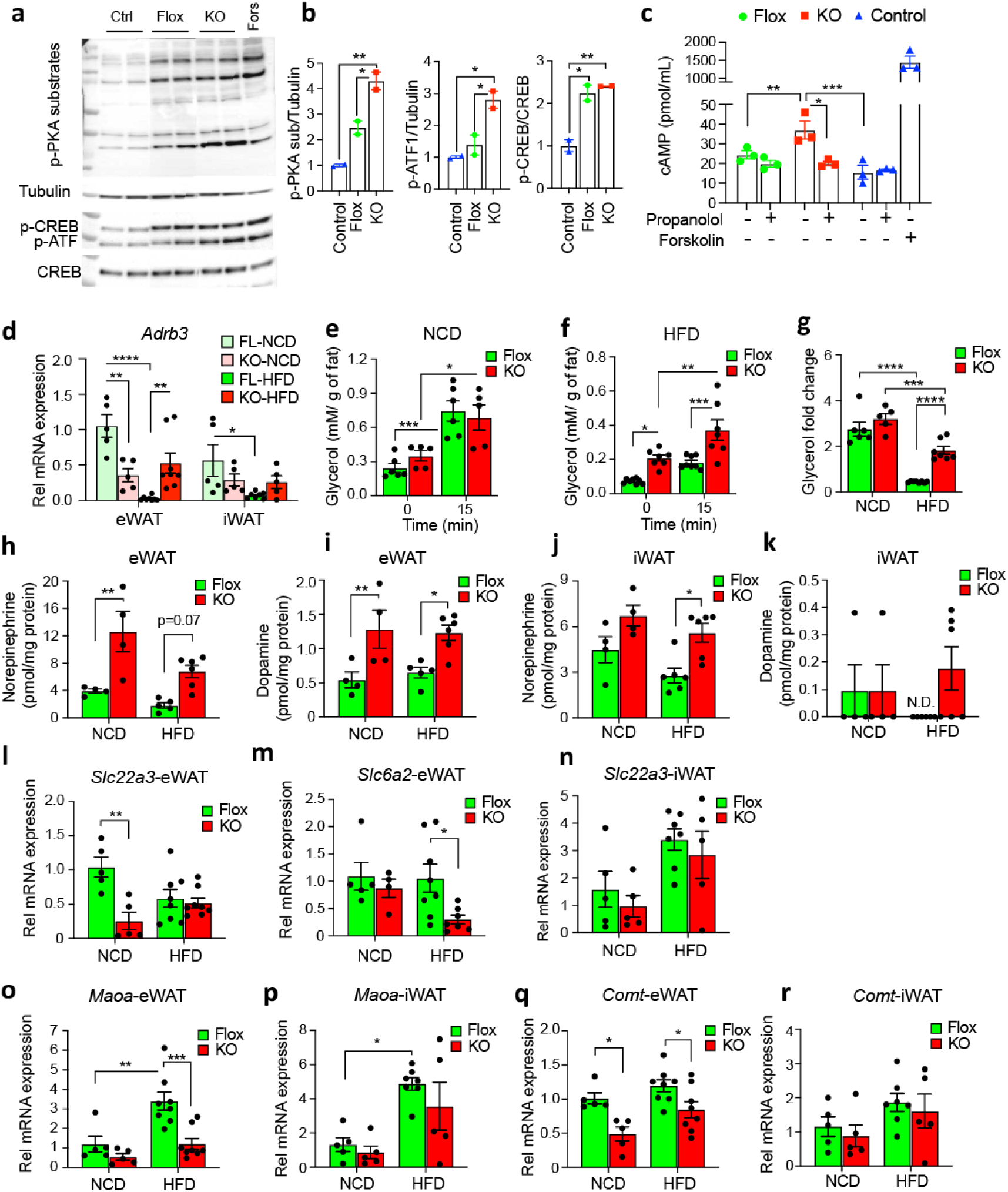
Norepinephrine and dopamine are elevated in WAT of Gnas^ΔCD11c^ KO mice. (**a**) Immunoblot for the phosphorylation of PKA substrates, and the phosphorylation of CREB and ATF1 in HEK293 cells treated for 30min with 3% serum from HFD-fed KO and Flox mice. Tubulin was used as loading control. (**b**) Quantification of p-PKA substrates, p-ATF1, and p-CREB in HEK 293 cells (n=2/group). (**c**) cAMP level in HEK293 cells treated with vehicle, forskolin (10 mM), 3% serum from KO or Flox mice, with or without the non-selective b-adrenergic receptor blocker propranolol (50 mM), by ELISA (n=3/group). (**d**) Relative mRNA levels of b3 adrenergic receptor (*Adrb3)* in eWAT and iWAT of NCD- or HFD-fed Flox and KO mice by q-PCR, normalized to m36B4 housekeeping gene (n=5-8 for eWAT groups, n=5- 7 for iWAT groups). (**e-g**) Serum glycerol levels before and 15 min after CL-316,243 injection in NCD-fed (**e**) or HFD-fed (**f**) Flox and KO mice per gram adipose tissue weight. (**g**) Fold increase in serum glycerol levels per gram of fat 15 min after CL-316,243 injection in the mice (n=5-8/group). (**h**-**k**) Norepinephrine and dopamine levels in eWAT and iWAT extracts of NCD-fed (n=4) or HFD-fed (n=6) Flox and KO mice determined by ultra-performance liquid chromatography (UPLC) normalized per mg protein. (**l-n**) Expression of the catecholamine transporter genes *Slc22a3* (OCT3) and *Slc6a2* (NET) in eWAT and iWAT of NCD- (n=5) or HFD-fed (n=8) Flox and KO mice by q-PCR normalized to m36B4. (**o-r**) Expression of catecholamine degradation enzymes monoamine-oxidase A (*Maoa)* and catecholamine-O-methyl transferase (*Comt)* in eWAT and iWAT. Data are represented as mean ± SEM, asterisks indicate statistical significance by one-way or 2-way ANOVA with Tukey’s multiple as appropriate; *p < 0.05, **p < 0.01, ***p<0.001, ****p < 0.0001.

Obesity desensitizes lipolytic signaling in white adipose tissue in response to b-adrenergic agonists^19^. To determine whether the KO mice are protected from catecholamine resistance, we gave a single intraperitoneal (i.p.) injection of the selective b3-adrenergic receptor agonist CL316,243 (CL) to the KO and Flox mice on either NCD or HFD and measured plasma glycerol levels. Injection of CL significantly increased the glycerol levels in the plasma of both KO and Flox mice on NCD despite the reduced b3AR mRNA in the KO WAT (**Fig. 6e**). Consistent with the greatly decreased b3AR in Flox mice on HFD, CL did not significantly increase glycerol levels in Flox mice (**Fig. 6f**). The KO mice had elevated glycerol levels both before and after CL treatment (**Fig. 6f**) suggesting increased lipolysis. When the data were expressed as the fold increase in plasma glycerol in response to CL treatment, the Flox mice showed catecholamine resistance on HFD with dramatically decreased induction of lipolysis, but the KO mice were relatively protected against catecholamine resistance (**Fig. 6g**).

The adrenergic receptors are receptors for the catecholamines norepinephrine (NE) and epinephrine (Epi) and at higher concentrations for dopamine (DA). Therefore, we measured catecholamine levels in NCD- and HFD-fed KO and Flox mice. We detected significantly higher NE and DA in eWAT and NE in iWAT of KO mice compared to Flox mice under both NCD and HFD feeding (**Fig. 6h-j**). DA was not detected in the majority of iWAT samples (**Fig. 6k**) and Epi was not detected in either eWAT or iWAT. Circulating and adrenal catecholamines were not altered between KO and Flox mice (**Supplemental Fig. 10c-h**). To explore the cause of the elevated catecholamines in the KO WAT, we measured expression of the catecholamine transporters and enzymes of catecholamine synthesis and degradation. Expression of *Slc22a3* was decreased in the eWAT of KO mice on NCD (**Fig. 6l**) but in contrast expression of *Slc6a2* was decreased in the eWAT of KO mice on HFD (**Fig. 6m**). Expression of *Slc22a3* did not change significantly in iWAT (**Fig. 6n**) and *Slc6a2* was not detected in iWAT. The expression of monoamine oxidase A (*Maoa)* was significantly decreased in both eWAT and iWAT from Flox mice with HFD, and this decrease was blunted in the KO mice (**Fig. 6o and p**). Expression of catecholamine-O-methyl transferase (*Comt*) was significantly lower in eWAT from KO mice (**Fig. 6q**) but did not change in iWAT (**Fig. 8r**). Expression of the rate-limiting enzyme for catecholamine synthesis, tyrosine hydroxylase (*Th*), did not differ in the adrenal glands of KO and Flox mice, and expression was not detected in the WAT of Flox and KO mice (**Supplemental Figure 10i**) suggesting that the elevated NE and DA in the KO eWAT most likely resulted from decreased catecholamine catabolism. Collectively, these data suggesting that Gas deficiency in CD11c+ cells promoted adipose tissue lipolysis through elevation of norepinephrine by downregulating catecholamine catabolism.

## DISCUSSION

We report in this paper that suppression of cAMP generation in CD11c+ cells by deletion of Gas caused both an increase in norepinephrine drive to adipose tissue and an increase in NE sensitivity that resulted in beigeing of adipocytes with increased lipolysis and thermogenesis (**Fig. 7**). The cumulative effects of these changes were to create a lean, obesity-resistant phenotype. The effect was mediated by bone-marrow derived cells and did not require adaptive immunity. Our results support the importance of the innate immune cell – adipocyte interaction in regulating adipose tissue function and provide novel insights into how sympathetic tone is regulated by the immune system at the tissue level.

**Figure 7:**
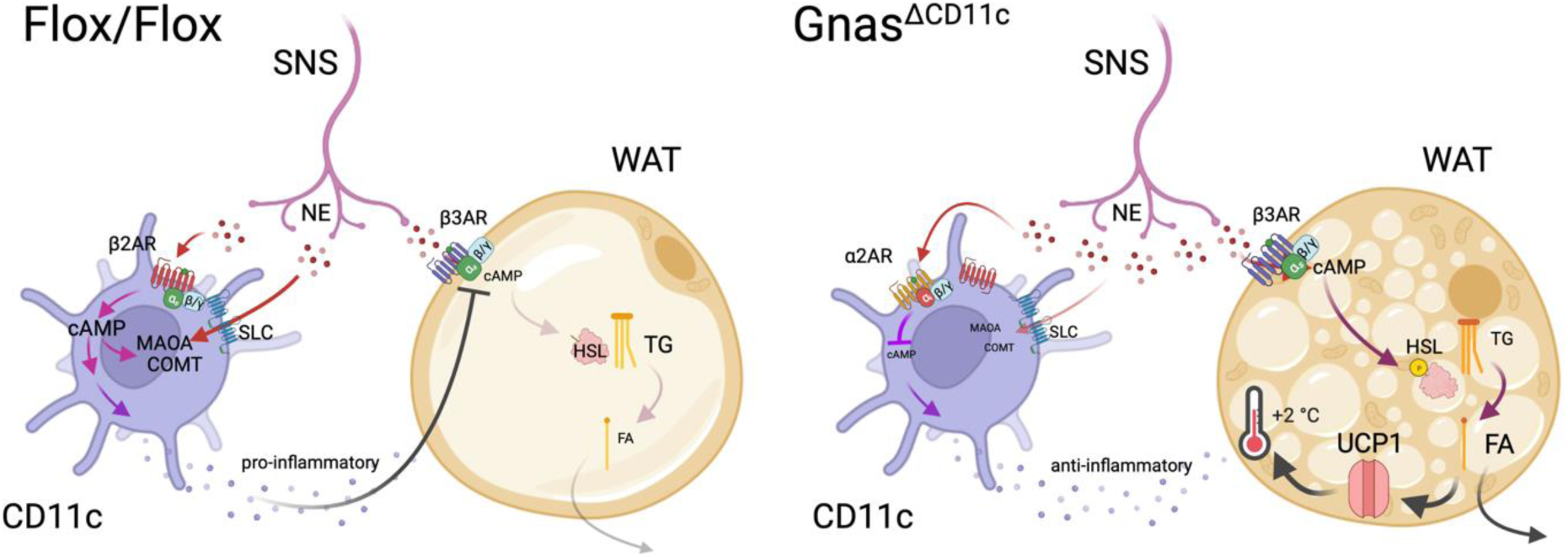
Schematic of crosstalk between adrenergic effects in CD11c immune cells and adipocytes. Norepinephrine (NE) stimulates CD11c+ cells via the b2-AR and Gas to elevate cAMP, which leads to an increase expression of MAOA and COMT and the production of pro-inflammatory cytokines. These cytokines act on adipocytes to inhibit b3-AR expression and cAMP generation to cause catecholamine resistance and limit lipolysis. In CD11c cells lacking Gas, the NE acts via a2-AR to inhibit cAMP generation which results in decreased MAOA and COMT expression and higher tissue catecholamine levels, and also the production of anti-inflammatory cytokines. The anti-inflammatory cytokines do not cause catecholamine resistance in the adipocytes, so the elevated NE activates cAMP signaling, increases lipolysis and liberation of fatty acids, and causes beigeing of the adipocytes with increased UCP1 expression and thermogenesis.

The connection between sympathetic activity and adipose tissue function has been studied in both animal models and humans^20–23^. Peripheral sympatho-facilitators, which cause an increase in sympathetic tone but avoid the adverse effects of global NE treatment^24, 25^, cause lipolysis, increased thermogenesis and weight loss without effects on food intake or activity^26^. Similarly, direct optogenetic stimulation of adipose tissue sympathetic nerves activates lipolysis and causes fat mass loss^11^. Complete loss of catecholamine production has no effect on DIO but mice have normal glucose tolerance^27^ suggesting that the ratio of b- to a-adrenergic signaling is important. Similarly, adipose tissue denervation eliminates the thermogenic response to a wide variety of stimuli including fasting and cold exposure^9^. Interestingly, BAT denervation causes compensatory beigeing of WAT, and iWAT denervation activates BAT^28^, suggesting crosstalk between the depots. Similarly, lipectomy increases SNS drive to BAT^29^ as does BAT transplantation^30^. NE inhibits pre-adipocyte proliferation as does a b1-specific agonist, but a b2-specific agonist increases preadipocyte proliferation^31^. b3AR-specific agonists rescue metabolic dysfunction in mice^32–34^, cause beigeing of iWAT more than eWAT^35^, and enhances lipolysis and insulin secretion^36^, but b2AR-specific antagonists improve glucose and insulin tolerance without affecting body weight^37^. The three bARs have been genetically deleted. b1AR KO are cold sensitive, have decreased BAT thermogenesis, and are more susceptible to DIO, glucose intolerance, and high TG^38^. The b2AR knockout mice are grossly normal but are leaner and have reduced adipose tissue^39^. b3AR KO mice are more susceptible to DIO possibly through increased a2AR signaling^40, 41^. Mice with deletion of all three bARs are cold-intolerant and obese with higher FFA and glycerol at basal and fasted states^42^ and bAR signaling is required for weight loss in response to a ketogenic diet^43^. Optogenetic induction of adipocyte uncoupling is sufficient to protect against DIO indicating that activation of thermogenesis is sufficient for the lean phenotype^44^.

The human data are less clear but generally agree with the animal data^45, 46^. Humans have a different distribution of BAT than rodents^47^ but BAT activation has been proposed as an anti-obesity therapy^45, 48^. Catecholamine reuptake inhibitors cause weight loss by increasing thermogenesis^49^ and high urinary catecholamine levels in patients with pheochromocytoma and paraganglioma are inversely associated with obesity^50^. Conversely, patients taking beta blockers have decreased energy expenditure and increased body weight^51^. The b3-agonist, Mirabegron, has recently been shown to improve glucose homeostasis in humans^52, 53^ but a side effect of b-agonist therapy is an increase in macrophage infiltration into adipose tissue that could worsen insulin resistance, and potential increases in blood pressure.

Although cAMP signaling was reduced in CD11c+ cells, cAMP signaling was elevated in adipocytes due to the increased sympathetic tone. Consistent with the lean phenotype that we observed, selective activation of cAMP signaling in adipocytes by chemogenetic activation of Gs increases energy expenditure, improves glucose metabolism and decreases fat mass^54^. Deletion of Gai2 in adipocytes similarly protects against DIO^55^. Conversely, genetic deletion of Gas in adipocytes reduces lipolysis and impairs thermogenesis, but the mice maintain normal body weight and energy balance as lipid synthesis is decreased^56^. The KO mice do not respond to b3-agonists, are more insulin sensitive, gain weight on HFD, and are cold intolerant due to a defect in brown adipocyte differentiation and low UCP1. In contrast, when the Gas is deleted in both adipocytes and immune cells, the phenotype is very different^57^. The KO mice are lean, have severely reduced adipose tissue, impaired glucose tolerance, and reduced thermogenic response to catecholamines or diet. The phenotype of these mice resembles that of our *Gnas^ΔCD11c^* KO suggesting that the loss of Gas in immune cells is dominant over the effect of Gas loss in adipocytes.

The nature of the CD11c+ cells that are responsible for our observed phenotype remains to be determined. Depletion of all CD11c+ cells has been achieved using bone marrow transplants from CD11c-DTR mice^7^. Diphtheria toxin treatment for 17 days acutely depletes CD11c+ cells after bone marrow reconstitution in the obese recipient mice. With this relatively short-term treatment, there is no weight loss, but glucose and insulin tolerance are normalized, however the interpretation may be more complicated as CD11c depletion induces CD64+Ly6c+ monocytes with increased capacity for TNF secretion. Recently, a population of sympathetic nerve associated CD11c+ macrophages (SAMs) have been identified that sequester and degrade norepinephrine in adipose tissue^58^. Whether these SAMs are responsible for our lean phenotype is unclear as deletion of Gas using LysM-cre did not have the same phenotype, but a limitation is that LysM-cre is not expressed in all macrophages^16^. These cells express both the NE transporter *Sl6a2* and monoamine oxidase A *Maoa;* and deleting *Slc6a2* in SAMs increased thermogenesis and prevented DIO similar to our knockout mice^58^. The role of the catecholamine transporters is controversial, however, as another group has published that a different transporter OCT3 (*Slc22a3*) is involved in NE uptake into adipocytes^59^, and the cytoplasmic form of the enzyme catecholamine-O-methyl transferase (S-COMT) is largely responsible for peripheral metabolism of catecholamines^60^. Interestingly, *Comt* is a target for PPARg and pioglitazone treatment increases COMT expression, reduces plasma catecholamines, and improves glucose tolerance in rats^61^ and COMT (Val158Met) genotype is associated with abdominal obesity and a predilection for unhealthy foods in humans^62, 63^. Inflammation and catecholamine metabolism have also been linked as activation of the NLRP3 inflammasome upregulates MAOA expression, and increases lipolysis in macrophages^64^, and decreases AT inflammation^65^.

A further finding from our study is that deletion of Gas in CD11c+ cells prevented HFD-induced catecholamine resistance in adipocytes. On the one hand, the b3-AR protein is internalized via b-arrestin-2 and the *barr2* KO mice are mildly protected from DIO in the absence of any change in norepinephrine^66^. On the other hand, b3AR gene expression is strongly suppressed by obesity, a b3-agonist, or insulin treatment^67–69^ possibly via increased Foxp1 expression that represses C/EBPb^70^. Catecholamine resistance is also created by inflammatory signaling leading to phosphorylation and activation of PDE3B^19^. The lack of downregulation of the b3AR in the *Gnas^DCD11c^* KO could be directly due to altered inflammatory signaling from immune cells to adipocytes, or indirectly due to the prevention of obesity and hyperinsulinemia. Activation of Gs signaling in CD11c BMDC causes production of inflammatory Th17 promoting cytokines such as IL-1b and IL-6 that could mediate the adipocyte catecholamine resistance, whereas suppression of Gs signaling increases production of Th2 promoting cytokines such as IL-4 and IL-10^15, 71^. The most striking change in plasma cytokines in our KO mice is the strong elevation of IL-10, especially on HFD where levels reach 200 pg/ml. The role for IL-10 in obesity is controversial however, as mice lacking IL-10 have increased thermogenesis^72, 73^ but AAV overexpression of IL10 or transgenic overexpression in muscle reverses obesity, and improves glucose and insulin resistance^74, 75^, and systemic IL-10 administration can ameliorate detrimental cardiac effects of HFD^76^. *Fas* mutant mice are protected from DIO and have elevated levels of IL-4, IL-10 and UCP1^77^. Results may be different in humans, however, as IL-10 has no effect on human adipocytes although it has anti-inflammatory effects on immune cells^78^.

In conclusion, we have identified a cAMP-dependent interaction between CD11c+ cells and adipocytes that regulates AT norepinephrine levels and catecholamine sensitivity resulting in a lean, obesity-resistant phenotype due to increased AT beigeing and thermogenesis.

## METHODS

### Mouse studies

C57BL/6 (B6) mice were purchased from Jackson Laboratories. *Gnas^DCD11c^* (*Gnas^tm5.1Lsw/tm5.1Lsw^/Tg(Itgax-cre)1-1Reiz*) mice and *Gnas^DLysM^* (*Gnas^tm5.1Lsw/tm5.1Lsw^/Lyz2^tm1(cre)Ifo^*) mice were generated as previously described^15^. Briefly, to generate *Gnas^DCD11c^* and *Gnas^DLysM^* mice, floxed *Gnas^fl/fl^* (*Gnas^tm5.1Lsw/tm5.1Lsw^*) mice were crossed with *CD11c-Cre* and *LysM-Cre (Lyz2-cre)* mice, respectively. Floxed mice were used as littermate controls. *Rag1-/-/Gnas^DCD11c^* (Rag.DCD11c) mice were generated by crossing *Rag1-/-* mice with *Gnas^DCD11c^* mice, and *Rag1-/-/Gnas^fl/fl^* (Rag.Flox) mice were used as their littermate controls. In metabolic balance studies, mice were placed in metabolic cages (Columbus Instruments) to assess their food intake, drinking, energy expenditure, O_2_ consumption, CO_2_ production and physical activity. Energy expenditure was calculated from VO_2_ and RER using the Lusk equation, EE in Kcal/hr = (3.815 + 1.232 X RER) X VO_2_ in ml/min. In some experiments, mice were fed high fat diet (HFD, D12492, 60% kcal from fat; 5.24 kcal/g, Research Diets Inc, New Brunswick, NJ) for 8-10 weeks to cause obesity, or subjected to cold room (4 ℃) stress for indicated time. Control mice were fed normal chow diet (NCD, rodent 5001). For the diet switch study, mice were fed with normal chow diet for 3 days and then switched to and fed with HFD for another 3 days. Mice used in this study were aged between 2 and 4 months unless otherwise stated. All mice were housed at 24 ± 2 ℃ on a 12 h light/12 h dark cycle (light on at 6 am) and the animals had free access to food and water prior to being placed in study groups. All procedures were approved by the Animal Care Committee of University of California San Diego, School of Medicine.

### Fecal microbiota transplantation

C57BL/6 germ-free female mice were obtained from UCSD Animal care program and maintained in flexible film isolators. Experimental mice were housed in Sentry SPP System (Allentown) on a 12 h light–dark cycle. Mice were fed with irradiated HFD or autoclaved 2019S (Envigo) and autoclaved distilled water ad libitum two weeks in advance fecal transplantation. Stool samples were collected freshly from HFD-fed KO and Flox female mice and suspended in PBS. Mice were gavaged with 100 μl of stool suspensions (0.1 g/mL) every week for 3 times. Body weight was measured every week using autoclaved beakers.

### Glucose and insulin tolerance tests

For glucose tolerance test (GTT), mice were fasted for 8 h and injected with D-glucose (2 g/kg) intraperitoneally. Blood glucose was measured by tail bleed at 0 min and monitored at intervals up to 120 min using a glucose meter (Easy Step Blood Glucose Monitoring System, Home Aide Diagnostics, Inc., Deerfield Beach, FL). For insulin tolerance test (ITT), mice were fasted for 6 h and injected with recombinant human insulin (0.4 U/kg) intraperitoneally. Blood glucose was measured at intervals up to 120 min. Terminal fasting glucose was also measured following 6 h fasting. Terminal fasting plasma insulin was measured using the mouse insulin kit (Meso Scale Diagnostics, Rockville, MD) and the homeostatic model of insulin resistance (HOMA-IR) was calculated as follows: (fasting serum insulin concentration (mU/ml)) x (fasting blood glucose levels (mg/dl))/(405)

### Core body temperature and infrared thermography

Implantable radio transmitters (Mini Mitter Respironics) were used to determine core body temperature and locomotor activity of freely moving Flox and KO mice. The mice were anesthetized using isoflurane, and transmitters were implanted into the peritoneal cavity. The animals were allowed to recover for 1 wk before baseline set recording for several days, and then core body temperature and locomotor activity were recorded over a 24 h period. For the cold challenge, cages were placed in the cold room at 4°C for 4.5 h and body temperature recorded. Cages were then returned to room temperature for 1 h.

Interscapular BAT temperature was measured using an infrared camera (FLIR Systems). The mice were anesthetized using isoflurane and placed in dorsal or ventral positions to acquire static thermal images at a focal length of 40 cm. Thermal images were acquired by infrared camera mounted on top of a tripod during the light phase. The maximum temperature in the interscapular area of mice was calculated using FLIR Tool software.

### Lipid absorption studies, lipid profiling and lipidomics

Mice were fasted overnight (≥16 hours) before lipid absorption experiments. sesame oil (400 mL/mouse) was administered by oral gavage, using a blunt ball-tipped syringe. Blood was collected by tail bleed at indicated time points, and plasma triglyceride was measured using triglyceride quantification colorimetric/fluorometric Kit (Sigma, MAK266).

For blood lipid profiling, 40 µL mouse blood samples collected from tail bleed were placed into the cholestech LDX cassettes within 8 minutes after collection and analysed using the cholestech LDX™ system for total cholesterol (TC), high-density lipoprotein cholesterol (HDL-C) and triglyceride (TRG) according to the manufacturer’s instructions. For the lipidomics analysis, blood was collected from mice fasted for 8 hours. A total of 50 µL serum was analyzed by the UCSD Lipidomics Core^79^.

To assess in vivo lipolysis, serum glycerol was measured in blood obtained from non-fasted mice before and 15 min after i.p. injection of CL316,243 (1 mg/kg). Serum glycerol were measured using glycerol assay kit (Sigma, MAK117) and normalized to the weight of body fat.

### Morphological studies

Mouse tissues were fixed in 10% neutral buffered formalin and embedded in paraffin. For histopathological evaluation, sections of 5-mm-thick were stained with haematoxylin and eosin according to standard protocols. For adipose tissue, pictures of representative areas from each section in ×100 magnification were taken, and Adiposoft software was used to calculate cell size of 3 images/section/mouse. Minimal 20 mm and maximum 100 mm thresholds were set for automated measurement of adipocyte diameter followed by manual correction. A frequency distribution was calculated for each group. Total adipocyte number within the distribution was subsequently calculated, and the frequency was converted to a percentage of total adipocytes counted.

### RNA isolation and real-time PCR

Total RNA was extracted from cells or tissues using an PureLink™ RNA Mini Kit (Thermo Fisher Scientific) in accordance with the manufacturer’s instructions. 0.5-1 µg RNA was transcribed to complementary DNA with the reverse transcription system (Invitrogen). Real-time PCR was performed on a Bio-Rad CFX384™ real-time PCR detection system using iTaq™ Universal SYBR® Green Super mix (BioRad). Primers used in this study were provided in **Supplementary Table 1**. Data were normalized to m36b4 and analyzed using the ^ΔΔ^CT method.

### Protein preparation and western blotting

Proteins from cells or tissues were prepared using RIPA buffer (ThermoFisher Scientific) containing protease and phosphatase inhibitors. Lysates were incubated for 30 min on ice and centrifuged for 15 min at 10,000 rpm at 4°C. Protein lysates were quantified using the micro-BCA protein assay kit (23225, ThermoFisher Scientific, New York, NY) using BSA as standard. Total protein lysates (10-20 μg) were separated on 10-15% polyacrylamide gels and transferred onto Polyvinylidene difluoride (PVDF) membrane (IPVH00010, Millipore Sigma). Membranes were blocked in 5% non-fat milk or BSA in PBS with 0.1% Tween 20 (PBST) for 1 h at RT and then incubated at 4°C overnight with the primary antibody at indicated concentration. Next the membranes were washed for 5 min X 4 times with PBST and incubated with HRP-conjugated secondary antibodies (Santa Cruz Biotechnology, SantaCruz, CA) for 1 hour. The members were visualized on X-ray films or ChemiDoc Imaging System (BioRad, Hercules, CA) following the reaction with the enhanced chemiluminescence substrate (SuperSignal™ West Pico PLUS Chemiluminescent Substrate, ThermoFisher Scientific). Actin, HSP90 or a-tubulin were used as the internal controls. The representative blotting bands were repeated in at least three mice. For western blotting: anti-phospho-PKA substrates (4056), anti-phospho-CREB (Ser133, 9198), anti-CREB (9104), anti-phospho-p70 S6 Kinase (Thr389, 9206), anti-p70 S6 (9202), anti-b actin (3700) antibodies were purchased from Cell Signaling Technology (Danvers, MA). Antibodies to UCP1 (ab10983), PGC1a (ab188102) were purchased from Abcam (Cambridge, MA). Antibodies were used according to manufacturer’s recommendation.

### Cell culture

The human embryonic kidney cell 293 (HEK293) cell line was obtained from ATCC and maintained in DMEM supplemented with 10% fetal bovine serum, 100 U/mL penicillin, and 100 μg/mL streptomycin at 37°C in an atmosphere of 5% CO_2_. Unless otherwise described, chemicals used in this study were all obtained from Sigma and dissolved in appropriate solvents.

### Hormones, cytokine measurements and ELISAs

Serum insulin was determined using a rat/mouse insulin ELISA kit (Millipore, Billerica, Merck KGaA, USA) as instructed. Serum adipsin (ab170244) and adiponectin (ab108785) were determined using a mouse adipsin/adiponectin ELISA kit (abcam). Cytokine/chemokine measurements in sera were performed using U-PLEX Adipokine assays (Meso Scale Discovery, Millipore).

For measurement of cAMP levels, HEK293T cells were serum-starved overnight and then treated with vehicle, forskolin (10 mM), 3% serum for 10 min at 37°C, in the presence or absence of the PDE inhibitor isobutylmethylxanthine (IBMX, 200 mM) and/or non-selective beta-adrenergic receptor blocker propranolol (50 mM). Cyclic AMP level was determined by cyclic AMP ELISA kit (Cayman, 581001) in HEK293T cells according to the manufacturer’s protocol.

### Flow cytometry

Epididymal fat pads were weighed, rinsed three times in PBS, and then minced and digested with 1 mg/ml collagenase type II (Worthington) in PBS supplemented with 2% fetal bovine serum (FBS) for 25 min at 37°C with shaking, followed by quenching with 10% FBS DMEM (Invitrogen). Cell suspensions were filtered through a 100-mm filter and centrifuged at 500 g for 5 minutes. Stromal vascular cell (SVCs) pellets were then incubated with RBC Lysis Buffer (eBioscience) for 5 minutes prior to centrifugation (500 g for 5 minutes) and resuspension in FACS buffer (2% FBS containing PBS).

SVCs were washed with FACS buffer and incubated with Fc Block for 20 minutes prior to staining with indicated fluorescently labeled primary antibodies or control IgGs, for 30 minutes on ice. After two times of wash, cells were used for analysis. Antibodies used for cell labeling were purchased from BD Pharmingen and eBiosciences. The data were acquired by a C6 Accuri flow cytometer (BD Biosciences) and analyzed by FlowJo Software.

### Bone Marrow Transplantation

The C57BL/6 male mice were randomly assigned into two groups (n = 8 mice/group) and were fed HFD for 8 weeks to induce obesity. To generate chimeric mice, the obese WT mice were lethally irradiated (950 Rads from a cesium source, split into 2 equal doses with a 4-hour interval to minimize gastrointestinal toxicity) and then reconstituted i.v. with 6x10^6^ bone marrow (BM) cells harvested from age- and sex-matched *Gnas^DCD11c^ or Gnas^fl/fl^* donor mice (i.e., *Gnas^DCD11c^* BM → *WT* recipients and *Gnas^fl/fl^* BM → *WT* recipients, respectively) the day following the irradiation. All the recipients received sulfamethoxazole/trimethoprim in drinking water for two weeks after irradiation. Six weeks after reconstitution, chimerism was confirmed by qPCR analysis of *Cre* genomic DNA expression in peripheral blood cells as we previously described^80, 81^. The mice were kept on HFD for another 13 weeks and the body weights were recorded weekly. GTTs and ITTs were carried out 7 weeks after BMT.

### Oxygen consumption rate measurements

For the Seahorse study, FACS-sorted splenic cDC2 (0.5x10^6^/well) and splenic CD4+ T cells (10^6^/well) of HFD-fed KO and flox mice were seeded into poly-D-lysine-coated XF96 plate and oxygen consumption rates (OCRs) were determined under basal conditions followed by the sequential addition of oligomycin (1 mM), FCCP (2 mM), and antimycin/rotenone (1 mM).

Oxygen consumption rates in fresh liver, soleus, epididymal fat pat, inguinal fat pat, brown adipose tissue were measured using the Oxygraph-2k (Oroboros Instruments, Innsbruck, Austria). Approximately 50-100 mg was transferred into ice-cold preservation medium (BIOPS solution). O_2_ fluxes in the indicated tissues were obtained upon CI substrates (glutamate/malate, L), ADP, followed by addition of CI substrate pyruvate (CI Ox), then CII substrate succinate (CI & CII Ox), then FCCP to uncouple mitochondria (mUC), then addition of rotenone to inhibit CI to assess CII uncoupled (CII ETS), and malonate/antimycin to inhibit CII and CIII activity to assess residual oxygen consumption (ROX).

### Determination of catecholamine contents by ultra-performance liquid chromatography (UPLC)

Catecholamine amounts were measured on a UPLC (Waters). The snap-frozen adrenal glands, eWAT, iWAT samples were thawed and homogenized in 300 mL of cold PBS and centrifuged at 4°C for 15 min at 15,000 x g. Plasma samples (200 µL) were thawed on ice and add 2 ng DHBA. Plasma, supernatants cleaned from residue, were collected and used for HPLC measurements. DA, NE and EPI concentrations were assayed using alumina extraction followed by separation and analysis with HPLC. Dihydroxybenzylamine (Sigma, 858781) was used as an internal standard in each tissue sample. The data were analyzed with Empower software (Waters).

### Statistics

Data were analyzed using Prism (Graphpad) and are presented as mean ± SEM or mean ± SD. Statistical significance was determined using the unpaired two-tailed Student’s t-test or 1-way or 2-way ANOVA followed by Tukey post-tests for multiple variables. A p value of < 0.05 was considered significant and is presented as *p < 0.05, **p < 0.01, ***p < 0.001, or ****p < 0.0001.

## ACKNOWLEDGEMENTS

This work was funded primarily by NIH grants R01HL141999, U01AI125860 and R01CA196853 to ER and NJGW, and in part by a VA Merit Review Award I01BX004848 and Senior Research Career Scientist Award IBX005224 to NJGW. The research was also supported by the NIH Cancer Center Support Grant P30CA023100, the Diabetes Research Center Grant P30DK063491, and the San Diego Digestive Disease Center Grant P30DK120515. We would like to acknowledge the assistance of Dr. Valeria Estrada and the Biorepository and Tissue Technology Shared Resource at the Moores’ Cancer Center.

## CONFLICT OF INTEREST

The authors have declared that no conflict of interest exists

---

**Supplemental Figure 1.**
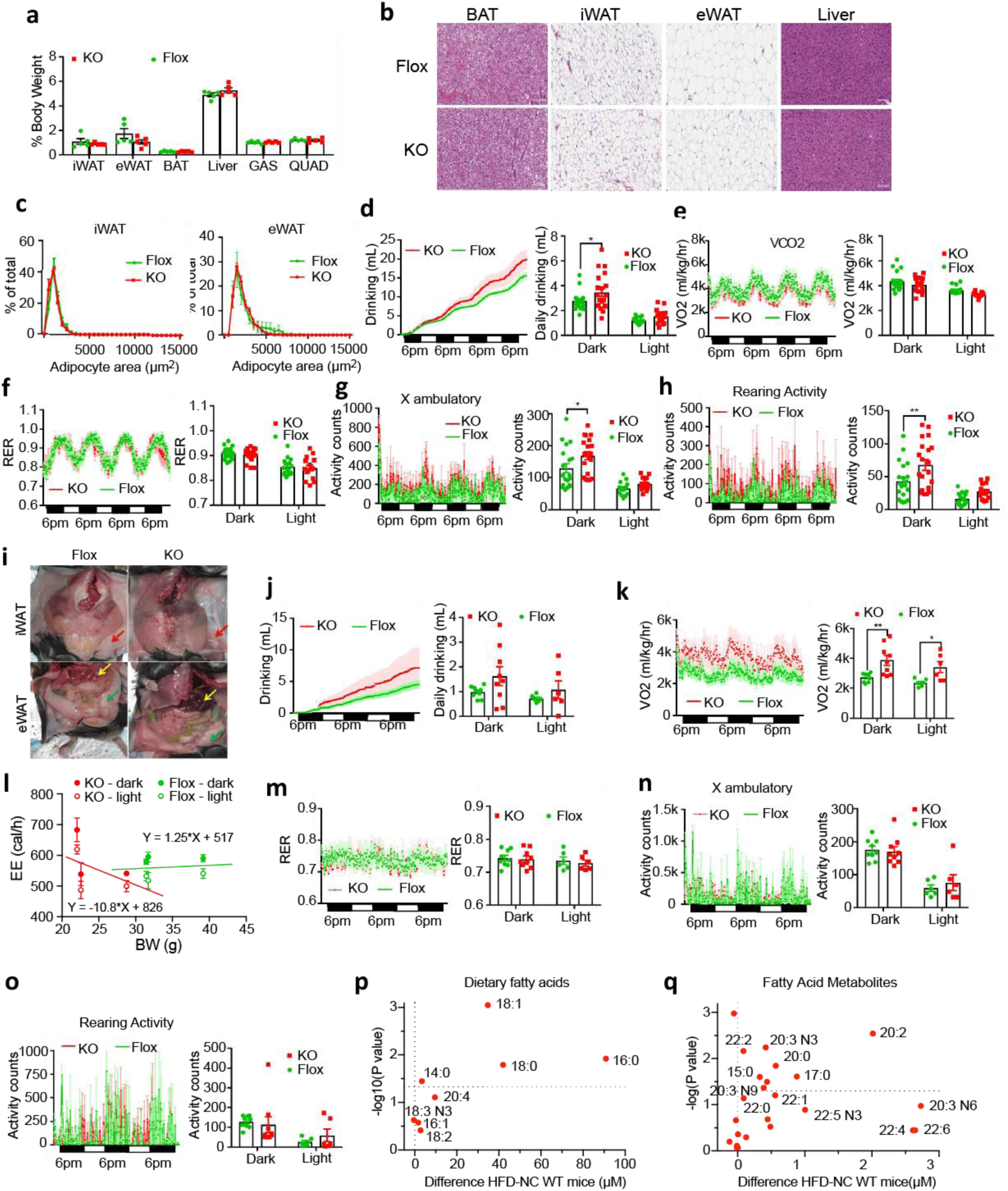
**(a)** Organ weights expressed as percent body weight (BW) of 4-month-old Flox and KO mice on NCD. Flox mice shown in green, KO mice in red. (**b**) Representative H&E staining of BAT, iWAT, eWAT and liver of NCD-fed Flox and KO mice, Scale bars=100 mm. (**c**) Adipocyte area distribution in iWAT (left) and eWAT from 4-mo-old male Flox and KO mice on NCD. n=5/group. (**d**-**h**) Assessment of energy expenditure in NCD-fed Flox and KO mice was performed using CLAMS. Black bars indicate the dark phase with lights out starting at 6 pm. (**d**) Cumulative drinking (left) and average drinking during dark or light cycle (right). (**e**) BW-adjusted carbon dioxide production rate (VCO_2_) over time and average VCO_2_ during light and dark cycle. (**f**) RER over time and average RER dark and light cycle. (**g**) Ambulatory counts over time and average ambulatory accounts during dark and light cycle. (**h**) Rearing activity counts over time and average rearing activity counts during dark and light cycles. (**i**) Representative images of eWAT and iWAT depots in Flox and KO mice on NCD and HFD. Red arrows indicate iWAT, green arrows eWAT and yellow arrows liver. (**j**-**n**) Assessment of energy expenditure in HFD-fed Flox and KO mice was performed using CLAMS; (**j**) Cumulative drinking and average drinking during dark or light cycle. (**k**) BW-adjusted VCO_2_ over time, and average VCO_2_ during light and dark cycles. (**l**) Absolute EE plotted against BW for KO (red) and Flox mice (green) for dark (solid symbol) and light (open symbol) phases with dependence derived from linear regression. (**m**) RER over time and average RER during dark and light cycle. (**n**) Ambulatory counts over time and average ambulatory accounts during dark and light cycles. (**o**) Rearing activity counts and average rearing activity counts during dark and light cycles. (**p**) Volcano plot showing plasma dietary fatty acid profiles for Flox mice on NCD and HFD. Plot shows the difference in FA concentration (µM) against the -log10 p-value. (**q**) Volcano plot for fatty acid metabolites for Flox mice on NCD and HFD. Plot shows the difference in metabolite concentration (µM) against the -log10 p-value. Data are represented as mean ± SEM and asterisks indicate statistical significance by 2-way repeated measures ANOVA or unpaired Student’s t-test as appropriate; *p < 0.05; **p < 0.01; ***p < 0.001.

**Supplemental Figure 2.**
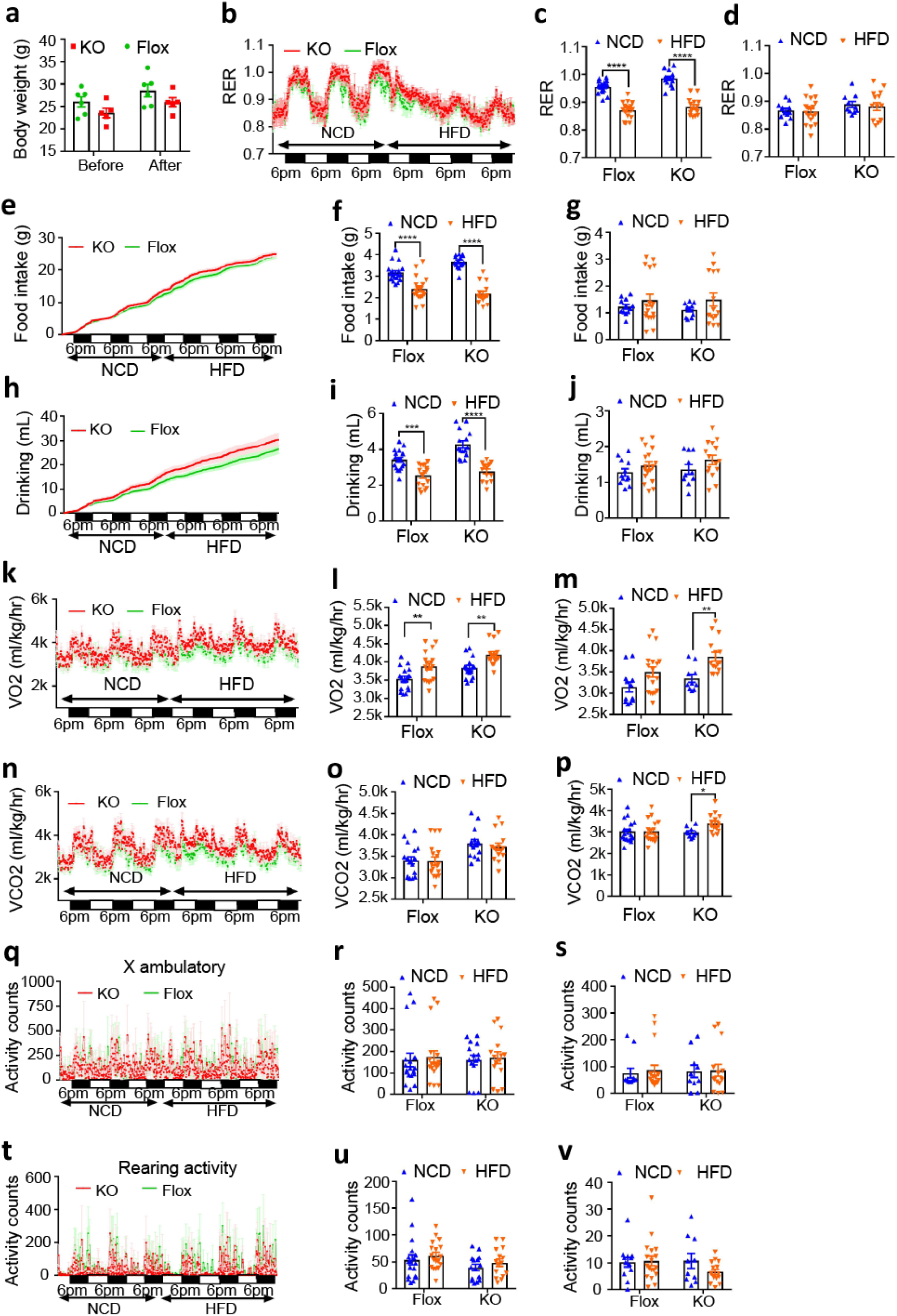
Food switch experiment. (**a**) Body weight of Flox and KO mice before and after the food switch experiment. n=6 for Flox group and n=5 for KO group. (**b**) RER over time and average RER during dark (**c**) and light cycles (**d**). Cumulative food intake (**e**), average food intake during dark cycle (**f**) and light cycle (**g**); cumulative drinking (**h**), average drinking during dark cycle (**i**) and light cycle (**j**); BW-adjusted VO_2_ (**k**), average VO_2_ during dark cycle (**l**) and light cycle (**m**); BW-adjusted VCO_2_ (**n**), average VCO_2_ during dark cycle (**o**) and light cycle (**p**); X ambulatory counts (**q**) and average counts during dark cycle (**r**) and light cycle (**s**); rearing activity counts (**t**) and average counts during dark cycle (**u**) and light cycle (**v**). Data are represented as mean ± SEM and asterisks indicate Statistical significance by 2-way ANOVA; *p < 0.05; **p < 0.01; ***p < 0.001; ****p<0.0001.

**Supplemental Figure 3.**
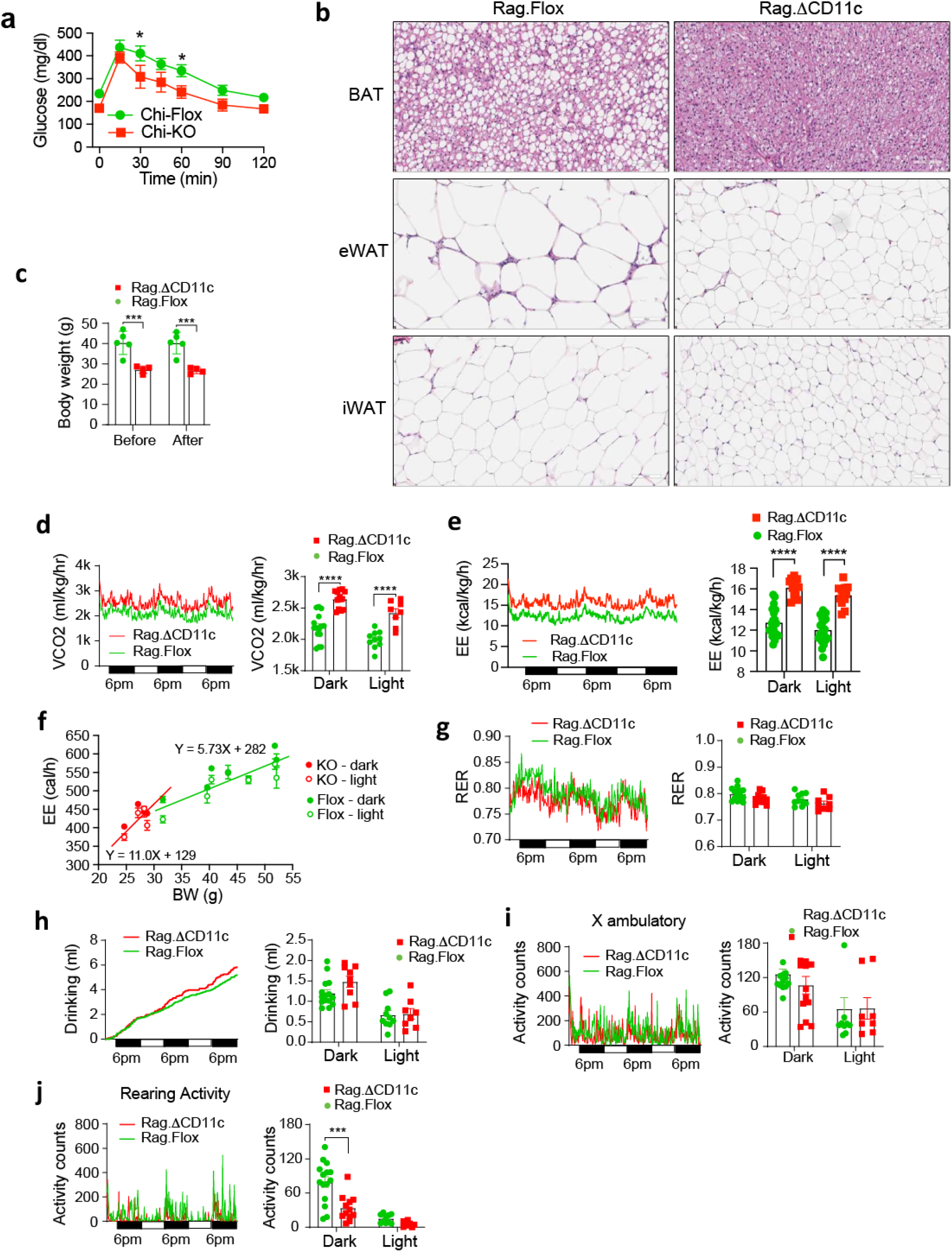
CLAMS study on HFD-fed Rag.DCD11c mice. (**a)** GTT for chimeric mice at week 11 after bone marrow transplantation. Chimeric mice that received KO bone marrow (Chi-KO) are shown in red, mice that received Flox bone marrow (Chi-Flox) in green. (**b)** Representative sections of BAT, eWAT and iWAT from Rag.Flox and Rag.DCD11c mice. (**c**) Body weights (BW) of the mice before and after the CLAMS study. Rag.Flox mice shown in green and Rag.DCD11c mice in red. (**d**-**j**) Assessment of energy expenditure in HFD-fed Rag.Flox and Rag.DCD11c mice was performed using CLAMS; (**d**) BW-adjusted VCO_2_ over time (left), average VCO_2_ during light and dark cycle (right). (**e**) BW-adjusted EE over time and average EE during light and dark cycle. (**f**) Absolute EE plotted against BW for KO (red) and Flox mice (green) for dark (solid symbol) and light (open symbol) phases with dependence derived from linear regression. (**d**) RER over time and average RER during dark and light cycle; (**e**) cumulative drinking and average drinking during dark and light cycle. (**f**) Ambulatory counts over time and average ambulatory accounts during dark and light cycle. (**g**) Rearing activity counts and average rearing activity counts during dark and light cycles. (**h**) Data are represented as mean ± SEM. n=5 for Rag.Flox group and n=4 for Rag.DCD11c group; ***p ≤ 0.001 and ****p ≤ 0.0001 by 2-way ANOVA.

**Supplemental Figure 4.**
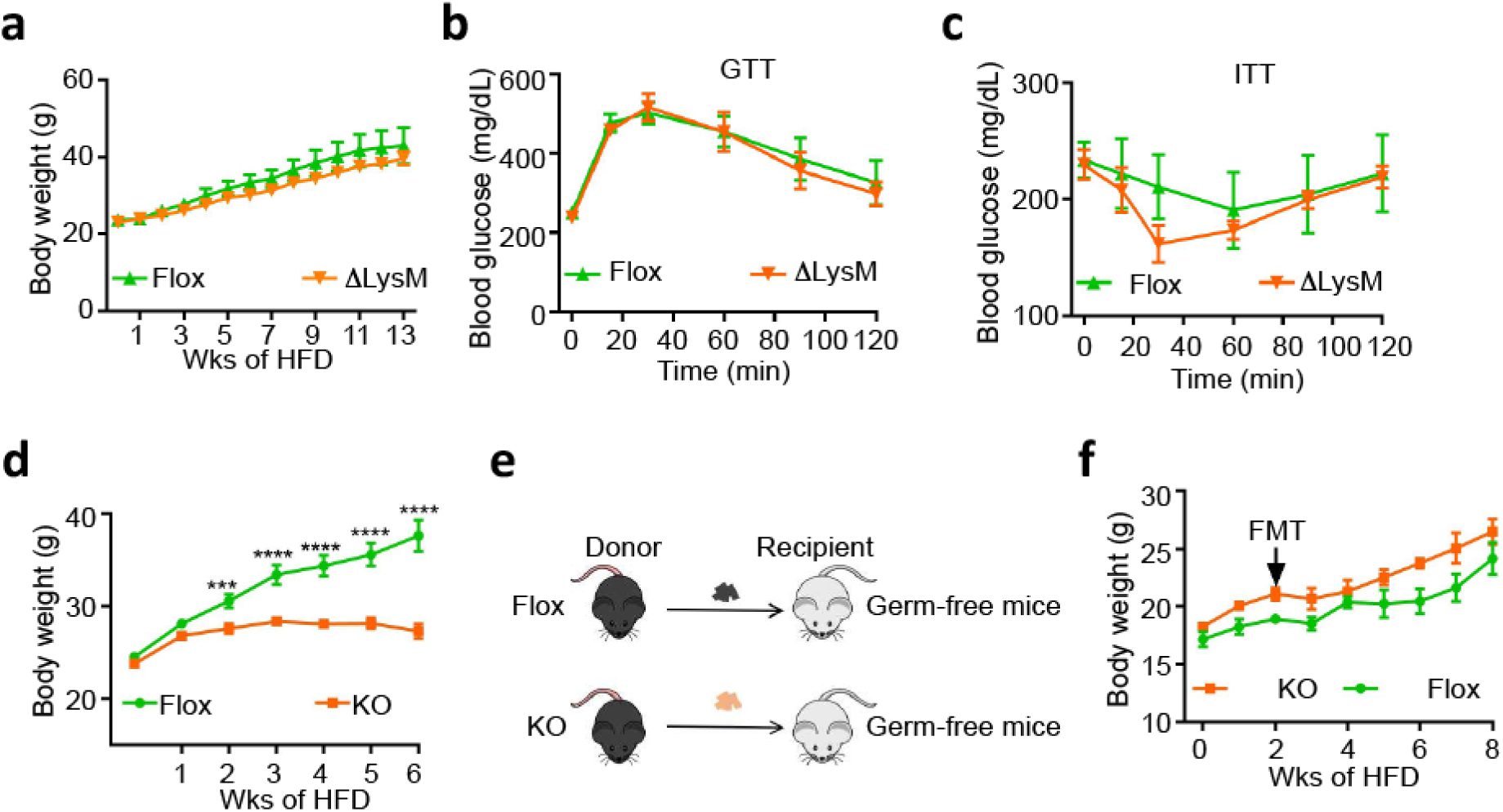
Deletion of Gnas in macrophages using LysM-cre and effect of microbiota on lean phenotype. (**a**) Body weights of Gnas^DLysM^ (DLysM) and Gnas^fl/fl^ (Flox) male mice on HFD were measured over time for 13 weeks. Glucose tolerance test (**b**) and insulin tolerance test (**c**) in Flox and Flox mice on HFD (n=6 per group). (**d**) Schematic diagram of fecal microbiota transplantation on germ-free mice. (**e**) 6 weeks-old germfree mice were first put on 60% HFD for 2 weeks. Thereafter, mice were orally treated with fecal microbiota of HFD-fed Gnas^fl/fl^ and Gnas^DCD11c^ mice weekly for 3 weeks and body weight were followed for another 6 weeks. n=4 per group. (**f**) The co-housed Gnas^fl/fl^ (Flox) and Gnas^DCD11c^ (KO) mice were put on HFD and body weights were recorded for 6 weeks. n=14 for Flox group, n=10 for KO group, ***p ≤ 0.001 and ****p ≤ 0.001 by 2-way ANOVA. Results represent mean ± SEM.

**Supplemental Figure 5.**
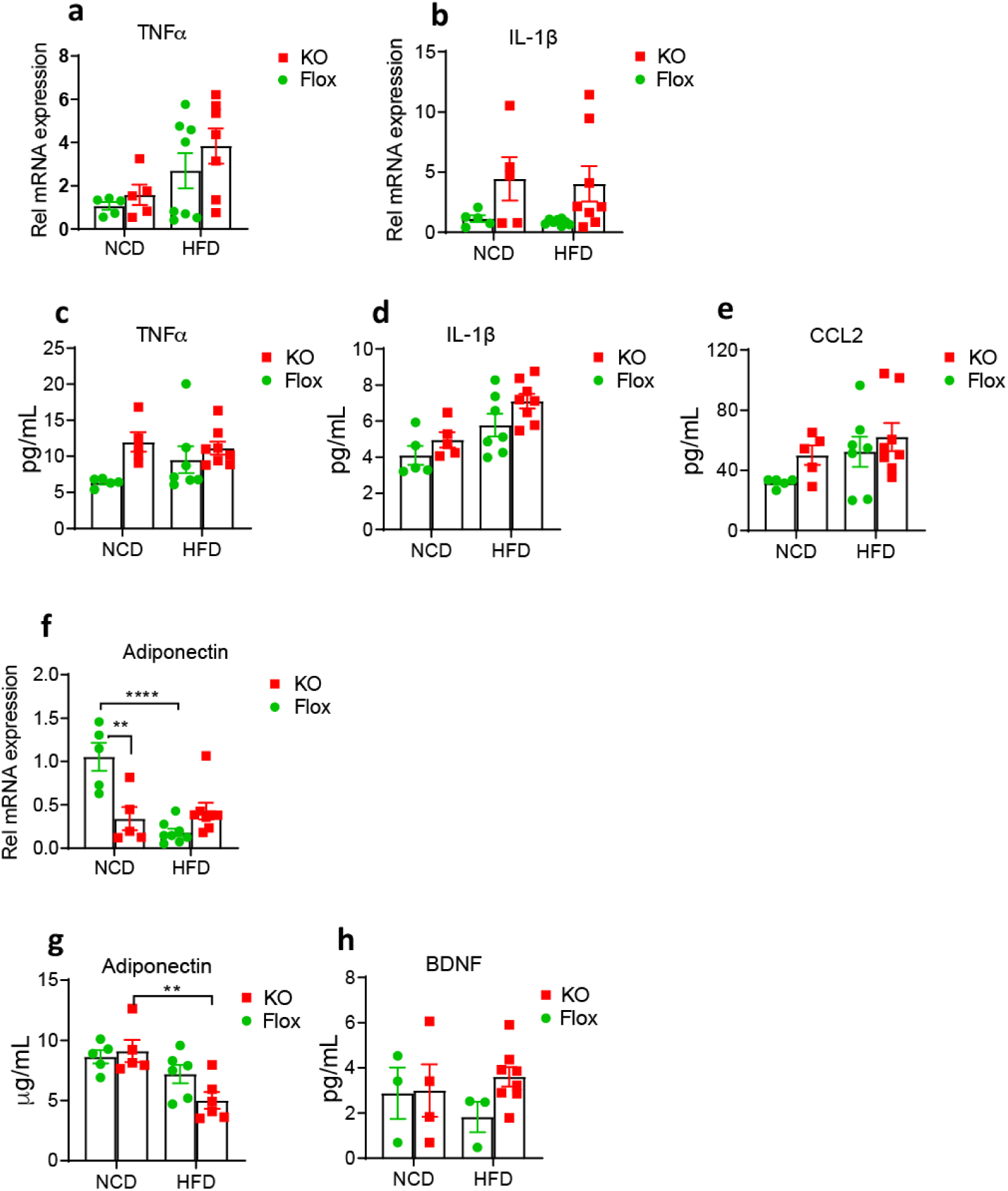
Cytokine and adipokine profiles of KO mice. Gene expression of inflammatory cytokine (**a**) TNFa, (**b**) IL1b and adipokine (**f**) adiponectin in the eWAT of KO and Flox mice fed with NCD or HFD for 8 weeks was analyzed by measuring relative mRNA levels using qPCR. The protein levels of (**c**) TNFa, (d) IL1b and CCL2, adipokine (**g**) adiponectin and (**h**) BDNF were measured in the plasma of Flox and KO mice. n=5-8 for each group, **p<0.01; **p<0.0001 by 2-way ANOVA. Data are represented as mean ± SEM.

**Supplemental Figure 6.**
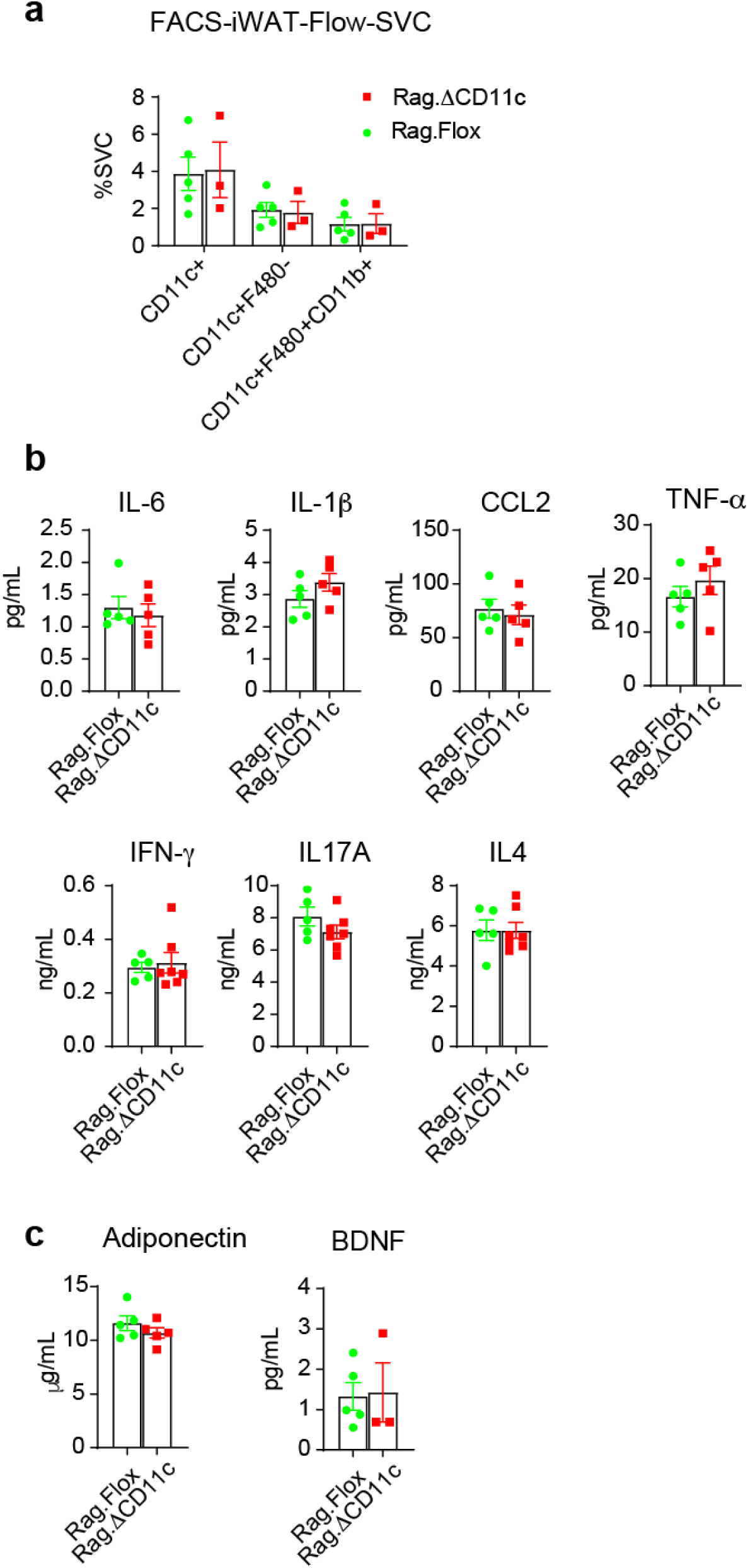
Cytokine and adipokine profiles of Rag.DCD11c KO mice. (**a**) SVCs were extracted from the iWAT samples and subsequently stained for F4/80, CD11b, and CD11c cell surface proteins and analyzed by FACS. n=4-5 per group. (**b**) The protein levels of cytokines IL6, IL1b, CCL2, TNF-a, IFN-g, IL17A and IL4 were measured in serum of HFD-fed Rag.Flox and Rag.DCD11c mice. n=5 per group. (**c**) The protein levels of adipokine Adiponectin, BNDF were measured in serum of the mice. Data are represented as mean ± SEM.

**Supplemental Figure 7.**
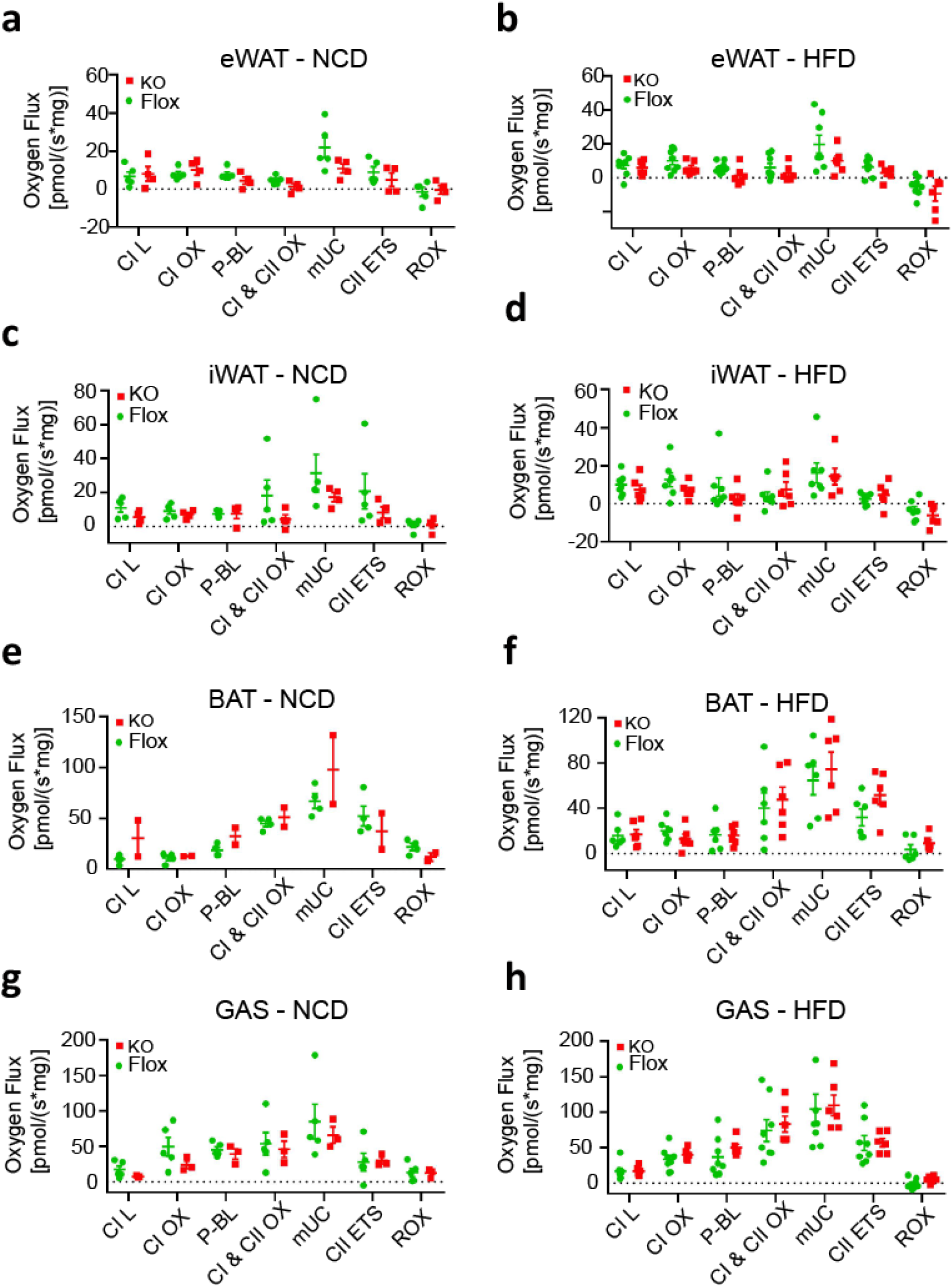
Oroboros oxygen consumption data in other tissues. O_2_ fluxes in eWAT (**a** and **b**), iWAT (**c** and **d**), BAT (**e** and **f**) and gastrocnemius (GAS, **g** and **h**) of Flox and KO mice with indicated diet upon CI substrates (glutamate/malate, L), oxidative phosphorylation (ADP added, Ox), followed by addition of CI substrate pyruvate (CI Ox), then CII substrate succinate (CI & CII Ox), then FCCP to uncouple mitochondria (mUC), then addition of rotenone to inhibit CI to assess CII uncoupled (CII ETS), and malonate/antimycin to inhibit CII and CIII activity to assess residual oxygen consumption (ROX). n=3-5 for NCD groups, n=6-8 for HFD groups. Data are represented as mean ± SEM.

**Supplemental Figure 8.**
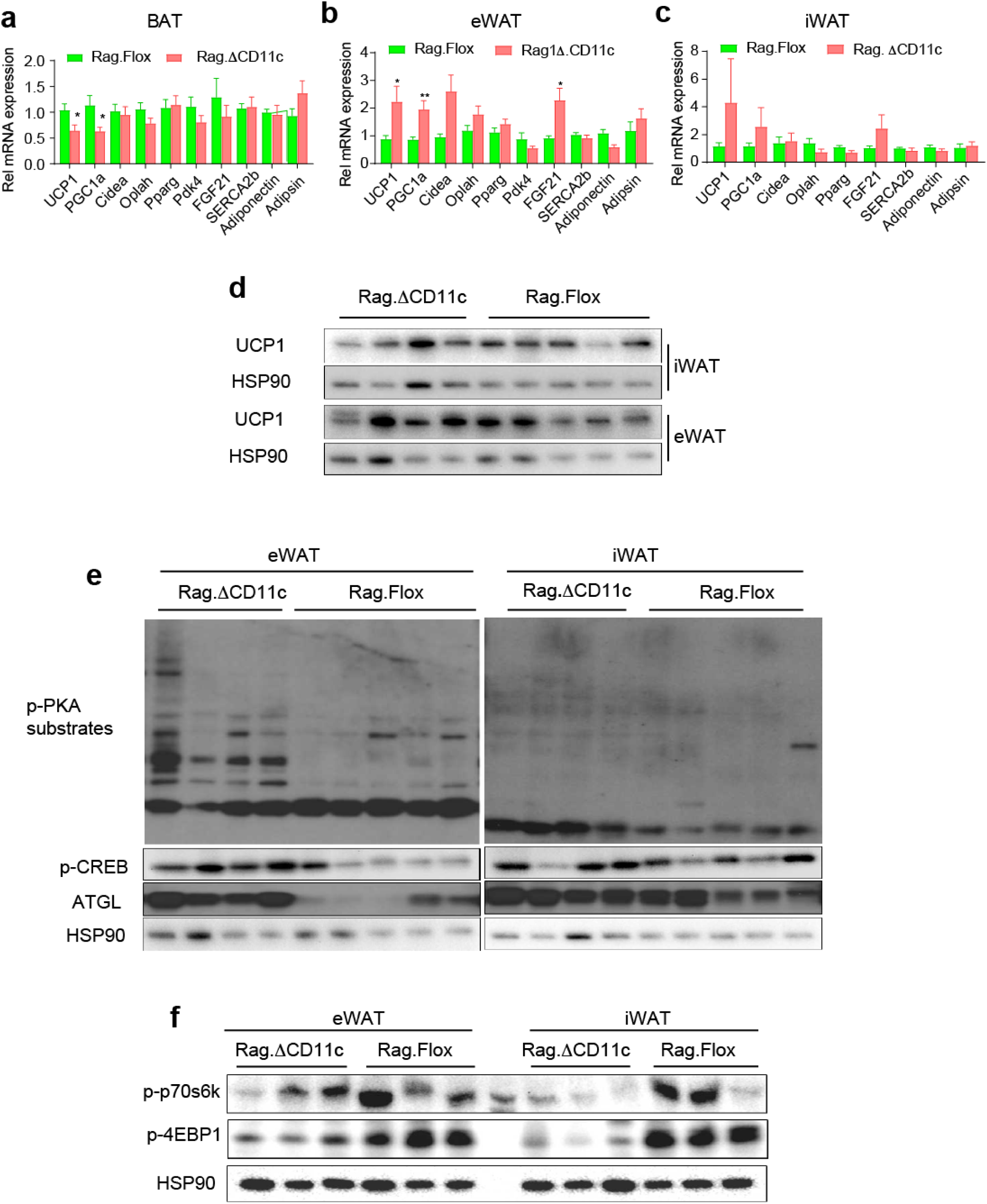
Increased adipose beigeing and PKA signaling in HFD-fed Rag.DCD11c KO mice. Relative mRNA levels of beigeing-associated genes in BAT (**a**), eWAT (**b**) and iWAT (**c**) of HFD-fed Rag.Flox and Rag.DCD11c mice, normalized to m36B4 control (n=6 per group). *p ≤ 0.05, **p ≤ 0.01 vs. control by Student’s t-test. (**d**) Representative Western blot of UCP1 in iWAT and eWAT of HFD-fed Rag.Flox and Rag.DCD11c mice, HSP90 was used as loading control. (**e**) Immunoblots of eWAT (left) and iWAT (right) from HFD-fed Rag.Flox and Rag.DCD11c mice probed with antibodies against p-PKA substrates, or antibodies against pCREB-S133, ATGL and HSP90. (**f**) Immunoblots of eWAT (left) and iWAT (right) probed with antibodies against pP70S6K-T389, p4EBP1-T37/46 and HSP90 (n=4-5/group). Data are represented as mean ± SEM.

**Supplemental Figure 9.**
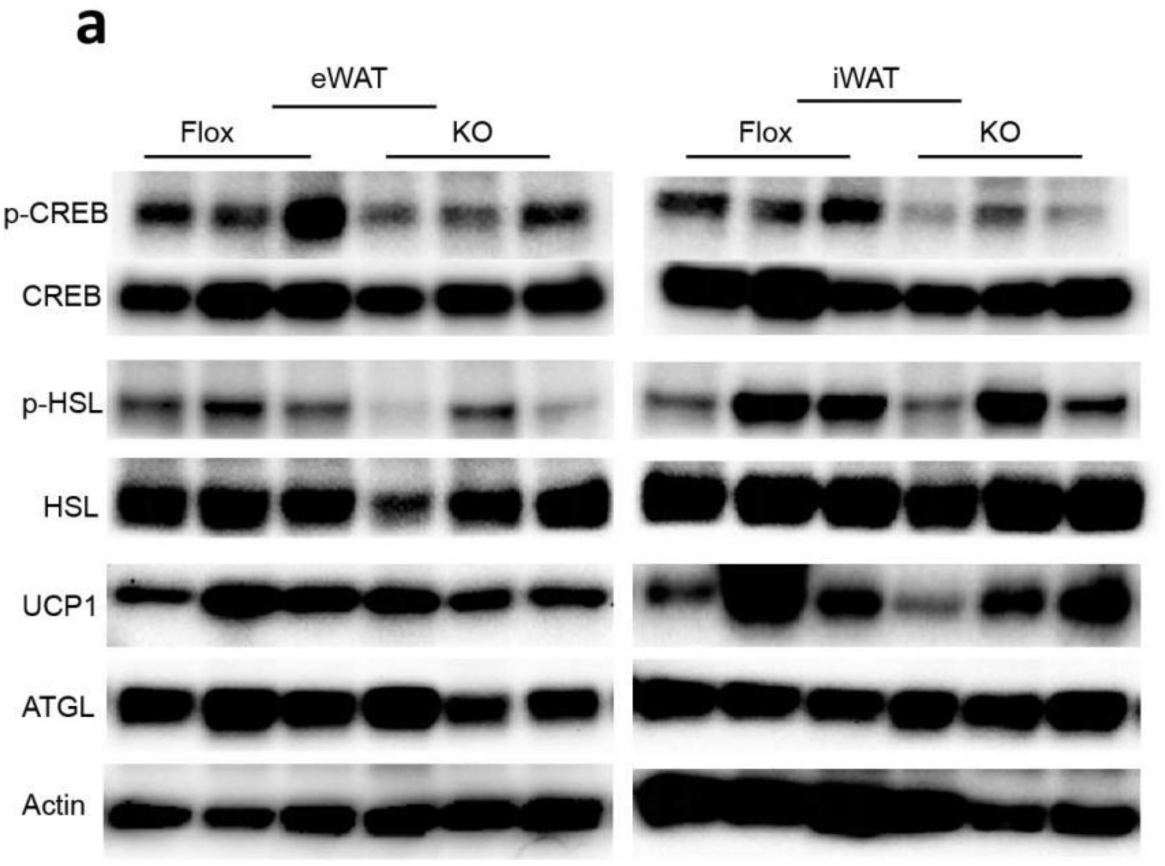
Representative Western blot of eWAT (left) and iWAT (right) from Flox and KO mice on NCD with antibodies against pCREB-S133/CREB, pHSL-S563/HSL, UCP1, ATGL and Actin. n=3 for each group.

**Supplemental Figure 10.**
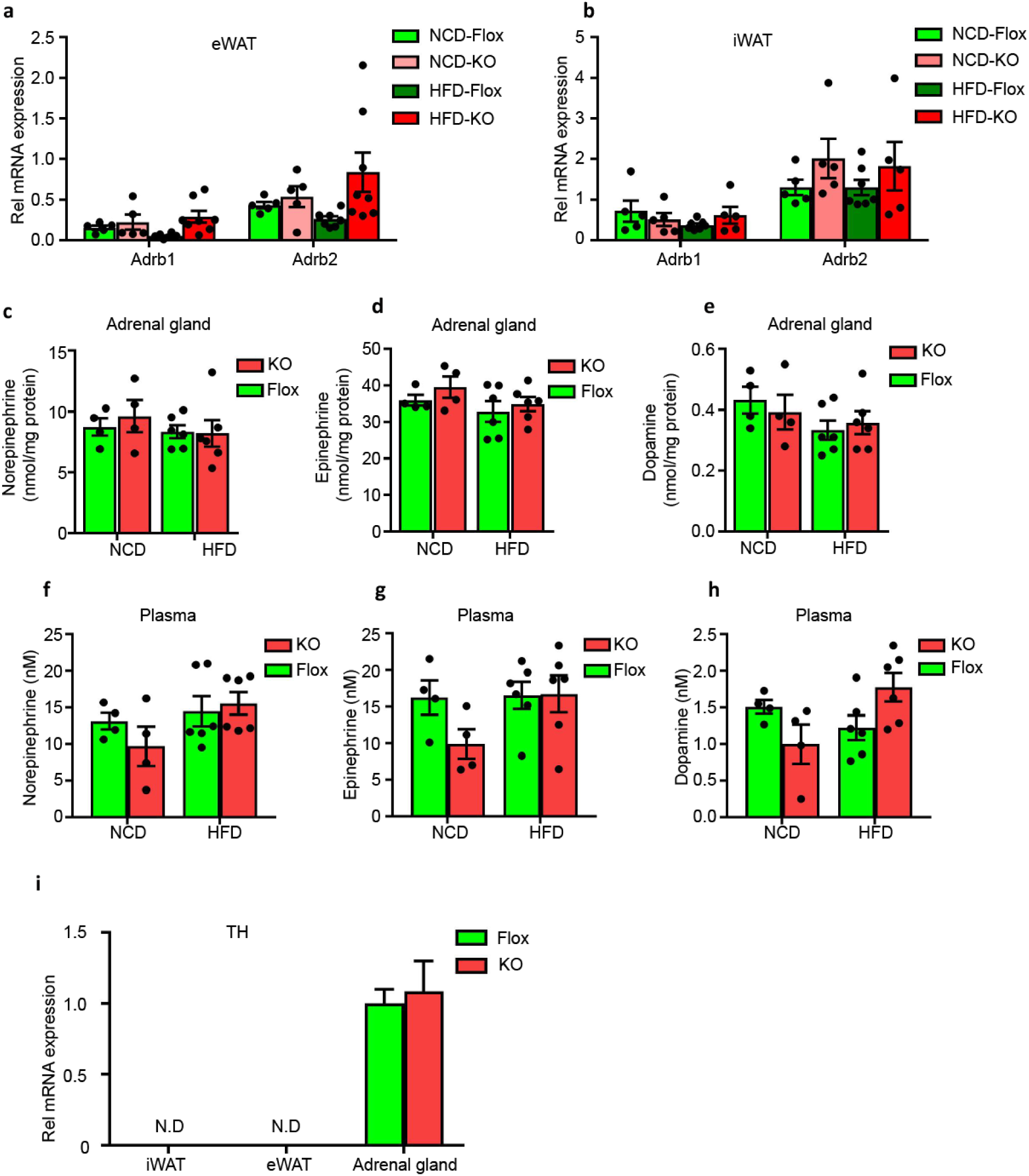
Adrenergic receptor expression and catecholamine levels in adrenal and plasma. Relative mRNA levels of beta-adrenergic receptors (Adrb1, 2) in eWATs (**a**) and iWATs (**b**) of NCD- or HFD-fed Flox and KO mice, normalized to m36B4 as housekeeping gene (n=5, 5, 8, 8 for eWAT groups, n=5, 5, 7, 5 for iWAT groups). (**c-e**) Adrenal norepinephrine, epinephrine and dopamine content by UPLC. (**g-i**) Plasma norepinephrine, epinephrine and dopamine by UPLC. (**j**) Expression of tyrosine hydroxylase (TH) mRNA was determined by qPCR normalized to m36B4 in iWAT, eWAT and adrenal glands of NCD- or HFD-fed Flox and KO mice. Data are represented as mean ± SEM.

## Notes

### Competing Interest Statement

The authors have declared no competing interest.

